# Biochemically distinct cohesin complexes mediate positioned loops between CTCF sites and dynamic loops within chromatin domains

**DOI:** 10.1101/2021.08.24.457555

**Authors:** Yu Liu, Job Dekker

**Affiliations:** Program in Systems Biology, Department of Biochemistry and Molecular Pharmacology, University of Massachusetts Medical School, Worcester, MA 01605-0103, USA; Howard Hughes Medical Institute, Chevy Chase, MD, USA

**Author notes:** Correspondence (J.D.).

## Abstract

The ring-like cohesin complex mediates sister chromatid cohesion by encircling pairs of sister chromatids. Cohesin also extrudes loops along chromatids. Whether the two activities involve similar mechanisms of DNA engagement is not known. We implemented an experimental approach based on isolated nuclei carrying engineered cleavable RAD21 proteins to precisely control cohesin ring integrity so that its role in chromatin looping could be studied under defined experimental conditions. This approach allowed us to identify cohesin complexes with distinct biochemical, and possibly structural properties, that mediate different sets of chromatin loops. When RAD21 is cleaved and the cohesin ring is opened, cohesin complexes at CTCF sites are released from DNA and loops at these elements are lost. In contrast, cohesin-dependent loops within chromatin domains and that are not anchored at CTCF sites are more resistant to RAD21 cleavage. The results show that the cohesin complex mediates loops in different ways depending on genomic context and suggests that it undergoes structural changes as it dynamically extrudes and encounters CTCF sites.

## INTRODUCTION

The cohesin complex plays major roles in mediating sister chromatid cohesion from S-phase to mitosis, and in folding chromosomes during interphase. The complex consists of two SMC proteins (SMC1 and SMC3) held together at their hinge domains, and a kleisin subunit (RAD21) that interacts with the SMC head domains. Additional subunits include SA1/SA2, NIPBL, and PDS5A/B^1–5^. Previous studies have provided strong support that the cohesin complex forms a ring-like structure that can entrap two sister chromatids^5–7^. During the metaphase-to-anaphase transition, separase cleaves RAD21 which opens the cohesin ring and releases sister chromatids so that they can be faithfully distributed to daughter cells^8–11^. Besides its critical role in sister chromatid cohesion, the cohesin complex is now understood to be an ATP-dependent motor protein that extrude chromatin loops all along chromatids throughout interphase^12–14^. Loop extrusion by cohesin has been observed using single molecule experiments^15–17^, and this activity requires the NIPBL subunit. Interestingly, CTCF-bound sites block cohesin-mediated loop extrusion in a CTCF site orientation-specific manner^18–23^, and this involves specific protein-protein interactions between a section of the N-terminal domain of CTCF and the SA2 and kleisin subunits of cohesin^24^.

Cohesin-mediated loop extrusion and directional blocking at CTCF sites produce chromosome structures that have been detected by Hi-C^12, 13^. First, stalled cohesins at CTCF sites produce relatively frequent and stable loops between pairs of convergent CTCF sites^18–21, 23, 25^. Such loops are visible as dots of elevated interactions in Hi-C contact maps. Second, cohesin mediates dynamic looping all along chromosomes, but rarely across CTCF sites. This leads to the formation of topologically associating domains (TADs) that are visible in Hi-C contact maps as domains with elevated intra-domain contact frequencies, and relatively sharp domain boundaries at CTCF sites. Interactions across such boundaries are relatively rare, due to the fact that they block loop extrusion, and the depletion of contacts across boundaries is referred to as insulation^26^. All these Hi-C features disappear when RAD21 is rapidly depleted from cells using an auxin-inducible degron approach^23, 25^: CTCF-CTCF loops, and domains of enriched interactions frequency (referred to here as TADs) disappear. On the other hand, other aspects of chromosome folding such as compartment domain formation, and compartmentalization are not lost.

A major open question is whether cohesin employs similar or different mechanisms to engage with chromatin when it mediates interactions between sister chromatids, maintains stalled loops between CTCF sites, or when it actively extrudes loops and travels along chromatin fibers within TADs. In an important earlier study, it was found that cleaving RAD21, which is sufficient to dissolve sister chromatid cohesion, has much milder effects on chromatin folding than complete degradation of RAD21 using a degron^23, 25, 27^. This suggests that different mechanisms may be at play for cohesion and extrusion respectively. Previous studies using RAD21 degron approaches were not able to address this question due to the complete loss of RAD21 and all cohesin-associated chromatin structures. Furthermore, as we show here, such degron approaches can suffer from confounding factors related to cell cycle changes that can obscure direct effects of the cohesin loss. Here we use a semi-in vitro approach using isolated nuclei expressing engineered cohesin subunits to study the effect of cohesin ring opening on chromosome folding in a controlled manner. Our results show that cohesin occurs in different biochemical, and possibly different conformational states when it mediates loops between CTCF sites, or when it mediates loops throughout TADs. Our data indicate that cohesin undergoes conformational changes and possibly subunit exchanges as it extrudes loops and then encounters CTCF sites.

## Results

### Effects of rapid degradation of RAD21 on chromatin folding can be obscured by cell cycle progression

Recent results have shown that degradation of RAD21 using auxin-inducible degron systems (Extended Data Fig. 1a and Fig. 1a) eliminates all loop and TAD structures in cells^23, 25^. However, we and others noticed that during auxin incubation cells continue to progress through the cell cycle and after 6 hours many cells are in G2/M phase (Fig. 1b)^28^. In mitotic cells, condensin complexes organize chromosomes to form rod-like structures, and TADs and CTCF-CTCF loops are completely lost^29–31^. Therefore, we were interested to determine the extent to which mitotic cells contribute to the observed changes in chromosome conformation upon RAD21 degradation (Fig. 1b). We used HCT116-RAD21-mAC cells to re-assess the effect of RAD21 depletion in G1 cells. We sorted and collected HCT116-RAD21-mAC G1 cells after RAD21 was degraded for 2 or 6 hours and performed Hi-C analysis (see methods). We also performed Hi-C analysis on nonsynchronized and G1/S synchronized (see Methods) HCT116-RAD21-mAC cells treated with IAA for 2 or 6 hours, as previously reported^25^. Examples of Hi-C maps are shown in Figure 1C. Visual inspection of the data shows that after 2 and 6 hours of IAA treatment, local features such as dots and domains become weaker or disappear (Fig. 1c and Extended Data Fig. 2a, cell cycle profiles in Extended Data Fig. 1b-d).

**Fig. 1.**
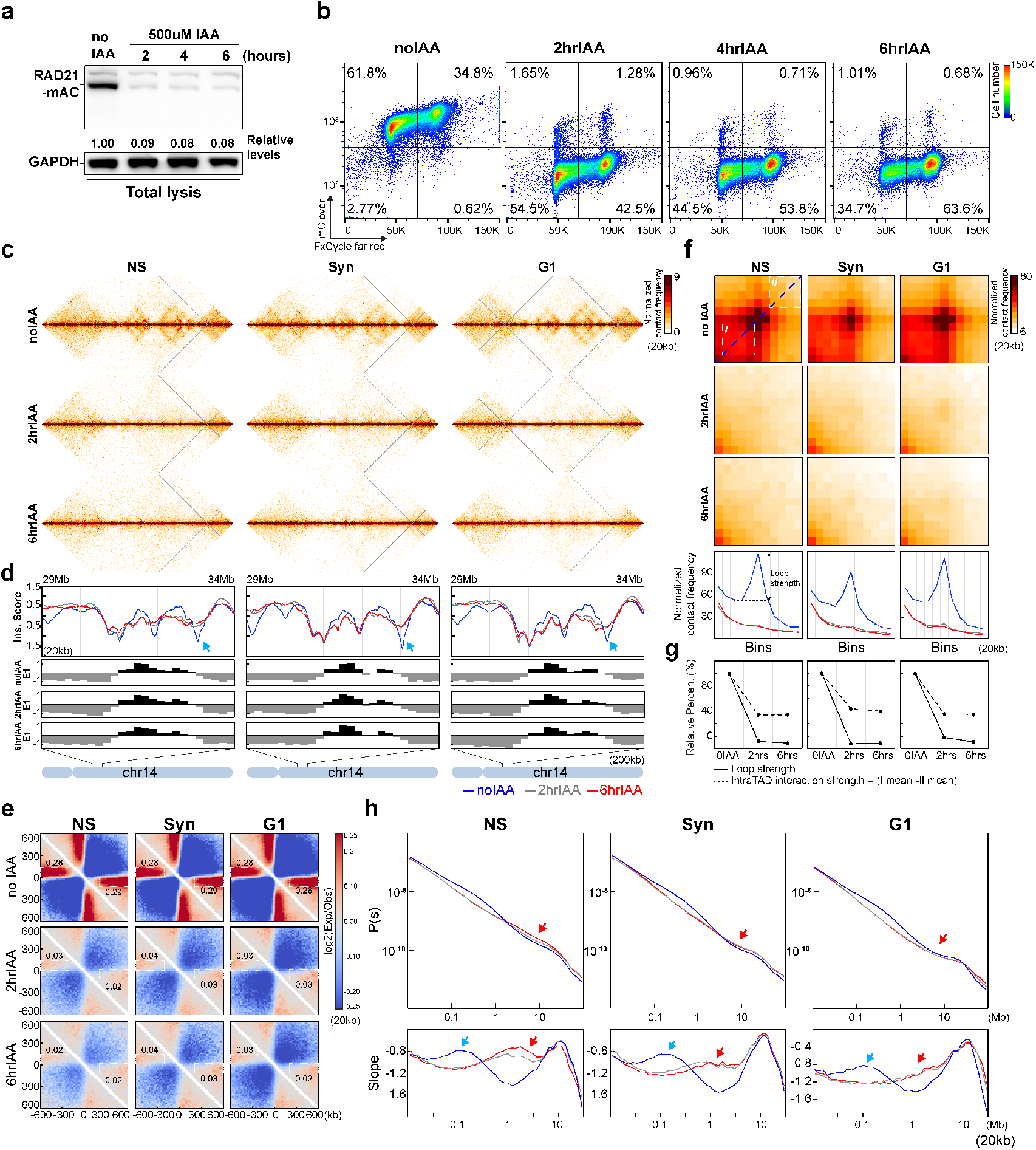
Degradation of RAD21 in G1 cells eliminates CTCF-CTCF loops and TADs. **(a)** Western blot analysis of RAD21 degradation in HCT116-RAD21-mAC cells after 500uM IAA treatments for 2, 4 and 6 hours as indicated. GAPDH was used as loading control. Numbers indicate the relative level of intact RAD21. **(b)** FACS analysis of HCT116-RAD21-mAC cells in absence of IAA and after 500uM IAA treatment for 2, 4 and 6 hours. DNA content was quantified using fxCycle-far red, and the level of RAD21 was quantified by mClover fluorence, The percentages in four squares indicate G1 cells with high RAD21 levels (top left), G1 cells with low RAD21 levels (bottom left), G2/M cells with high RAD21 levels (top right), and G2/M cells with low RAD21 levels (bottom right). **(c)** Hi-C interaction maps for non-synchronous (NS), synchronous (Syn), and G1 cells without and with IAA treatment for 2 and 6 hours respectively. Data for the 29-34 Mb region of chromosome 14 are shown at 20kb resolution. **(d)** Insulation profiles for the 29-34 Mb region of chromosome 14 at 20kb resolution. The blue, grey and red lines represent no IAA treatments, 2 hour IAA and 6 hour IAA treatments, respectively, as shown. The panels F and H have the same color assignments for the lines as in panel D. The lower panels indicate compartment Eigenvector value E1 across the same region at 200kb resolution. **(e)** Aggregate Hi-C data binned at 20kb resolution at TAD boundaries that were identified in each condition of NS, Syn and G1 without IAA treatments. The number of TAD boundaries for NS, Syn and G1 are 4227, 4272 and 4210, respectively. The numbers at the sides of the cross indicate the strength of boundary-anchored stripes using the mean values of interaction frequency within the white dashed boxes. These values are related to boundary strength. **(f)** Aggregated Hi-C data at a set of loops identified in HCT116-RAD21-mAC cells with intact RAD21 (n=3169) identified by^25^. Plots at the bottom show average Hi-C signals along the dotted blue lines representing signals from the bottom-left corner to the top-right corner of the loop aggregated heatmaps shown in upper panels **(g)** Quantification of loop strength and intra-TAD interaction strength. Loop strength was measured as the relative peak height of the plots in panel F. Relative peak height was calculated as the signal difference between the central and 3^rd^ bins (as indicated in the left bottom panels of **(f)**. Intra-TAD interaction strength represents relative contact frequencies within TADs, calculated as the average signal difference between interactions within region I within TADs and the region II outside TADs. Regions I and II are highlighted as white dashed squares in upper left heat map of panel F. **(h)** *P(s)* plots (upper panels) and the derivative from *P(s)* plots (lower panels) for Hi-C obtained from cells grown as indicated. The red arrows indicate the increased interaction frequency at s=10Mb (upper panels) and the appearance of condensin loop arrays structures (lower panels). The blue arrows indicate the signature of cohesin loops in each condition without IAA treatment. NS and Syn experiments were repeated once as a confirmation of previous studies. Hi-C experiments for G1 cells were repeated independently for three times and Hi-C results of G1 cells in Figures 1 and 2 are from the pool of three replicates.

To quantify loss of domanial features such as TADs we calculated insulation profiles genome-wide. Domain boundaries display local minima in insulation^26^. We observed the weakening and loss of many boundaries upon RAD21 degradation (Fig. 1d upper panel, black arrows; and Extended Data Fig. 2b black arrows), whereas compartment profiles remained unaffected (Fig. 1d, lower panel). To quantify loss of domain boundaries we first identified boundaries using the insulation profiles (see Methods). In each condition we discovered over 4,000 boundaries. We then aggregated Hi-C interactions at boundaries. In untreated cells we found the typical Hi-C features associated with TAD boundaries: strong depletion of interactions across boundaries, and line-like features of increased interactions anchored at the boundaries (Fig. 1e). After 2 or 6 hours of RAD21 depletion both these features were strongly diminished. This was observed for all three conditions (non-synchronized cells, synchronized cells and G1-sorted cells (Fig. 1e and Extended Data Fig. 2c).

We then examined the effect of RAD21 depletion on loop formation. We aggregated our Hi-C data at a set of 3,169 loops, mostly CTCF-CTCF loops and referred to as such from here on, identified by Rao et al in HCT116 cells^25^. In untreated cells we readily detect the strong focal enrichment of interactions between the loop bases (Fig. 1f, Extended Data Fig. 2e). After 2 or 6 hours of IAA treatment, this focal enrichment is completely lost (Fig. 1e and Extended Data Fig. 2d) in non-synchronized, synchronized and G1 sorted cells. To quantify loop strength, we plot the Hi-C data along the diagonal line of loop aggregation heatmaps (Fig. 1f, the blue dashed line on the heatmap of non-synchronized (NS) with no IAA). The resulting plots reveal peaks at the position of the looping interaction (blue lines in the plots at the bottom of Fig. 1f). We use the relative height of these peaks (as indicated in the plots) as a measure for loop strength. We observed that loop strength is reduced to below zero after RAD21 depletion, in all conditions. The negative loop strength is the result of the general distance-dependent decay in interaction frequency in the absence of looping interactions.

Extruding cohesin also mediates randomly positioned loops within TADs, resulting in elevated contact frequencies within TADs as compared to interactions between TADs. To quantify enrichment of intra-TAD interactions, we calculated the average intra-TAD interaction frequency (indicated by white square number I in the top right panel of Fig. 1f) and the average interaction between loci outside the domain (indicated by white square number II in the top right panel of Fig. 1f). We then calculated the difference between these averages and use this as a metric for intra-TAD interaction strength. In untreated cells interactions within the TAD are strongly enriched, and this is greatly reduced upon RAD21 depletion (Fig. 1g). The remaining intra-TAD strength (~35% of the initial strength; Fig. 1g) represents the base line level in the absence of TADs and is due to the known general distance-dependent decay in interaction frequency: interactions within white square I are for loci that are closer together in the linear genome as compared to interactions within white square II. We conclude that RAD21 depletion leads to loss of all positioned loops (at CTCF sites) as well as all extruding loops within TADs.

Finally, we calculated how interaction frequency (*P*) decayed with genomic distance (*s*). In untreated cells we observe the typical shape of *P*(*s*), with a relative shoulder where interactions decay more slowly for loci separated by ~100 kb (Fig. 1h, top plots). The derivative of *P*(*s*) highlights this shoulder as a local peak (Fig. 1h bottom plots blue arrows). Previous work has established that this shoulder, and corresponding global peak in the derivative of *P*(*s*), represents the average size of loops^32^. Upon RAD21 depletion this shoulder, and peak in the derivative of *P*(*s*), disappears entirely (Fig. 1h) in all three conditions reflecting the loss of cohesin-mediated loops^33^. However, we observed that in non-synchronized, and to some extent in synchronized cells, interactions between loci separated by 1-2 Mb increased (Fig. h, upper plots, red arrows). This is observed in the derivative of *P*(*s*) where a new peak is observed at around 1 Mb (Fig. 1h lower plots, red arrows). This is reminiscent of what is observed in mitotic cells. In mitotic cells condensins generate loop arrays and in human cells this is apparent in the *P*(*s*) plots by a shoulder at ~1 Mb^29, 33^. In G1-sorted cells we did not detect this phenomenon. Therefore, we believe that in non-synchronized, and to some extent in synchronized cells, at least a sub-population of cells entered mitosis during RAD21 depletion. This is consistent with data shown in Figure 1b. Therefore, under those conditions the presence of mitotic cells in the cultures can contribute to the loss of CTCF-CTCF loops and TADs. In G1-sorted cells this confounding factor is not present, making the interpretation and quantification of effects of RAD21 depletion more accurate.

RAD21 degradation did not lead to significant changes of compartment boundaries as determined by the main E1 eigenvector (Fig. 2a-b). Interestingly E1 values for B compartment domains became more negative in G1-sorted cells after RAD21 depletion (Fig. 2b, arrow heads). To quantify compartmentalization strength, i.e., the preference of compartment domains to interact with other domains of the same type, we calculated saddle plots ^22^. In these plots, loci are ranked by their E1 value and pairwise interactions are calculated and normalized for their expected interaction frequency given their genomic distance. Compartment strength is then calculated as the ratio of AA and BB to AB and BA interactions (see Methods). A slight increase of compartment strength was observed in non-synchronized cells after RAD21 was degraded for 2 hours (Fig. 2c, the left and middle columns of panels), whereas compartment strength decreased after RAD21 was degraded for 6 hours for non-synchronized cells (Fig. 2c). In both synchronized and G1-sorted cells, compartment strength increased after RAD21 depletion, with the largest increase in G1-sorted cells (Fig. 2c, the right column and Figure S2H for replicate experiments). The increased compartmentalization strength is consistent with previous studies that found that knocking out of the cohesin loading factor, NIBPL increases compartmentalization ^34^.

**Fig. 2.**
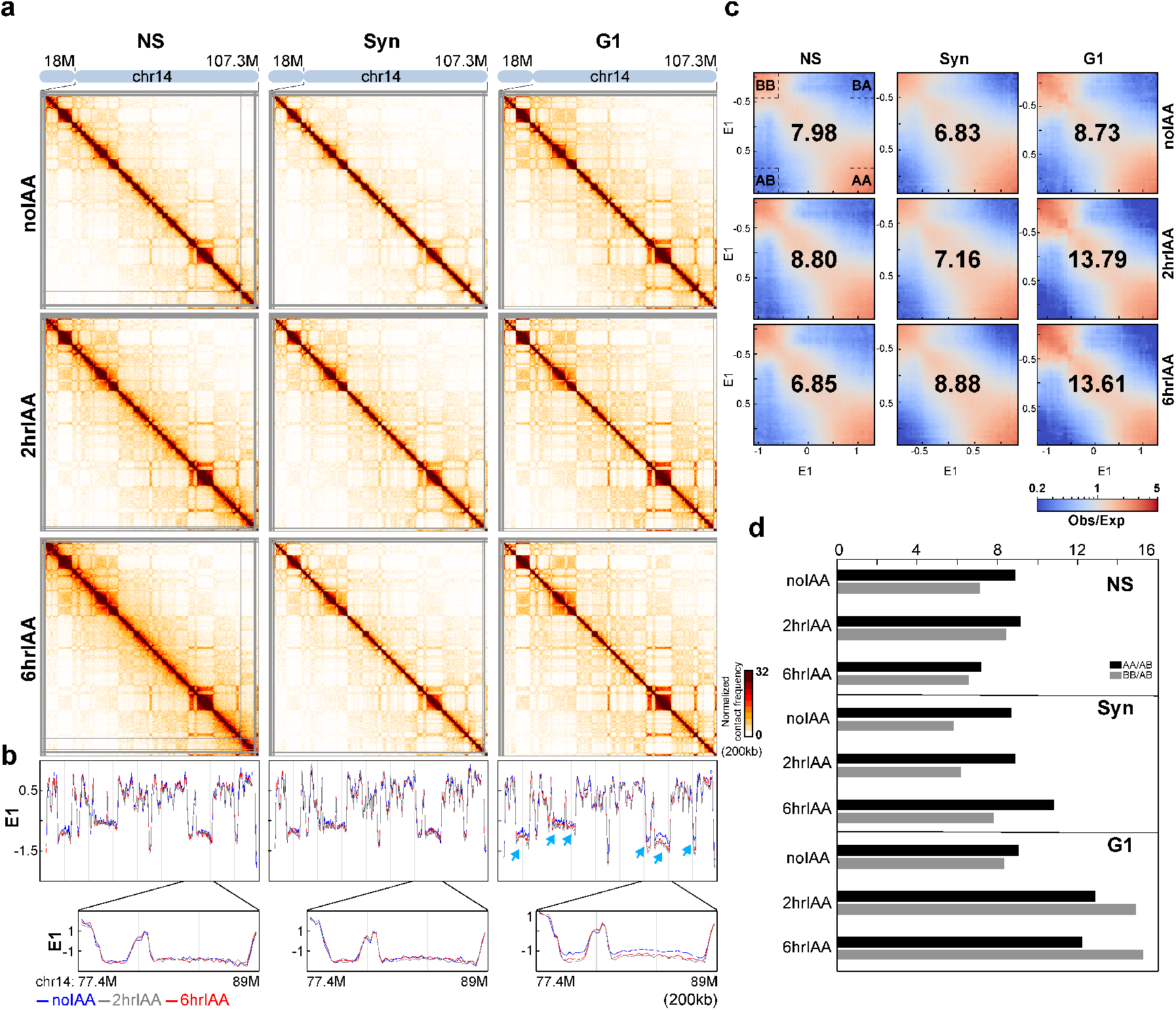
Degradation of RAD21 in G1 cells enhances compartmentalization. **(a)** Hi-C interaction maps for NS, Syn, and G1 cells without and with IAA treatment for 2 and 6 hours respectively. Data for the 18-107.3 Mb regions of chromosome 14 are shown at 200kb resolution. **(b)** Eigenvector value E1 across the 18-107.3 Mb region of chromosome 14 at 200kb resolution. The blue, grey and red lines represent: no IAA treatments, 2hour IAA and 6hour IAA treatments, respectively. Bottom, E1 for the 77.4-89Mb is shown at 200kb resolution. Blue arrows indicate changes of E1. **(c)** Saddle plots of Hi-C data binned at 200kb for NS, Syn and G1 cells without and with IAA treatments for 2 and 6 hours. Saddle plots for each cell condition were calculated using the E1 calculated from the Hi-C data obtained with cells grown without IAA treatments. The numbers at the center of the saddle plots indicate compartment strength calculated as the ratio of (A-A + B-B)/(A-B + B-A) using the mean values from the corners indicated by dashed boxes. **(d)**. Interaction strength of compartments. Dark bars indicate strength of A-A interaction as compared to AB interaction (A-A/A-B) while grey bars represent strength of BB interactions as compared to BA interaction (B-B/B-A).

We then examined the effects of RAD21 degradation on the strength of A-A and B-B interactions separately. In G1 cells, loss of RAD21 led to greatly increased compartmentalization for both A-A and B-B interactions, with B-B interactions becoming the strongest (Fig. 2d). In synchronized cells the effect on compartmentalization is more modest. In non-synchronized cells both A and B compartmentalization strength is reduced after 6 hours of RAD21 depletion, and preferential B-B interactions are the weakest. One explanation for the quantitatively different effect of RAD21 depletion in non-synchronized cells is that in those cell cultures many cells progress through the cell cycle and arrest in G2/M, as shown in (Fig. 1b and 1h). Mitotic cells do not display compartmentalization ^30^. Therefore, the effect of RAD21 depletion on compartmentalization is obscured by the fact that cells progress through the cell cycle during the auxin treatment.

In synchronized cells, compartment strength increased more slowly as compared to G1-sorted cells, Possibly, the chemicals used to synchronize cells in G1 affect compartmentalization dynamics. We note that the loop strength in synchronized cells was relatively weak even in cells without IAA treatment (Fig. 1f, the first row of heatmaps).

Taken together our results show that RAD21 degradation eliminated CTCF-CTCF loops and TADs, while it significantly enhanced compartment strength in G1 cells. In addition, our analysis shows that cell cycle dynamics, and unintended effects of protocols to arrest cells need to be taken into consideration when performing and interpreting depletion experiments.

### A system for controlled cohesin perturbation

We set out to establish an alternative biochemical experimental system for analysis of effects of cohesin perturbation on chromosome folding. Our approach uses purified nuclei that can be treated in various ways e.g., incubated in different buffer conditions etc. Previously, we have shown that chromosome folding in purified nuclei is comparable to that in intact cells^35^. An advantage of nuclei is that they do not progress through the cell cycle during the experiments, avoiding any of the confounding factors described above.

To allow for controlled perturbation of RAD21, we used CRISPR/Cas9 to edit the RAD21 gene in HAP1 cells so that they express a RAD21 protein that contains three repeats of the TEV protease recognition motif (HAP1-RAD21^TEV^ cells). The TEV motifs were inserted between Pro^471^ and Pro^472^ in exon 11 of RAD21 (Fig. 3a and Extended Data Fig. 3a). The TEV motifs were inserted in an unstructured region of RAD21 that connects the N- and C-terminal domains of the protein and their insertion is expected to have no effects on four key regions of RAD21 that interact with SMC1, SMC3, NIBPL and STAG1 (Fig. 3a)^36^. In previous work, TEV sites had been introduced in the corresponding position in *Scc1*, the mouse homolog of RAD21 (Extended Data Fig. 3a) and it was shown that the protein was fully functional^9^. HAP1-RAD21^TEV^ cells proliferate normally. The unstructured region in which the TEV sites were inserted contains one of two naturally occurring separase cleavage sites ^8^. Previous work has shown that cleaving RAD21 (or Scc1) by separase, or by TEV, in this region suffices to disrupt sister chromatid cohesion^8–11^. Furthermore, Zuin and co-workers showed that cleavage of RAD21 at another location within the unstructured region that connects the N- and C-termini of the protein leads to reduced TAD boundary formation, although the effects were not as strong as those observed after RAD21 depletion using IAA-inducible degron approaches^27^.

**Fig. 3.**
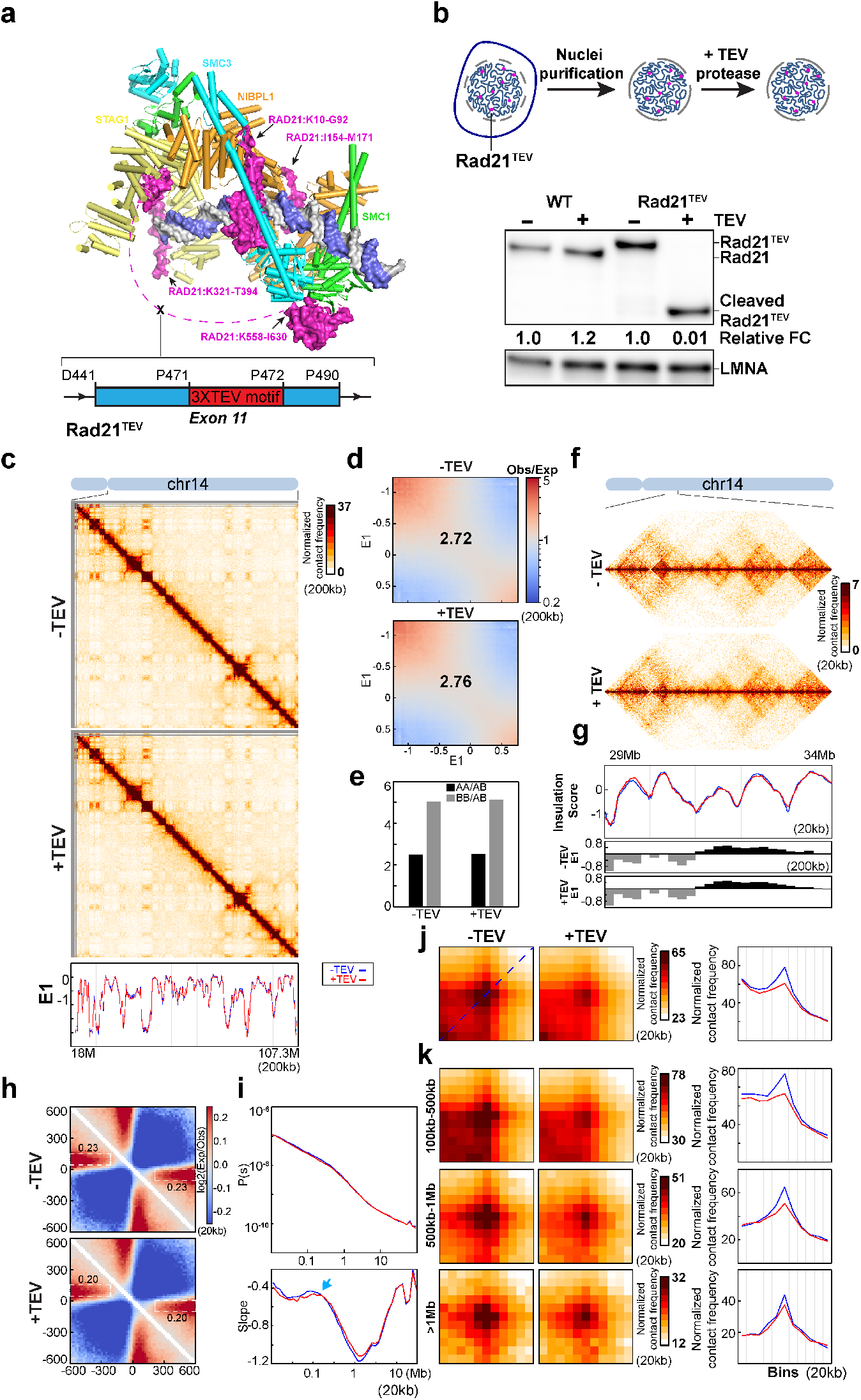
Cleaving RAD21 reduces CTCF-CTCF loop interactions. **(a)** Structure of the cohesin complex in association with DNA. The structure was drawn using 6WG3 from Protein Data Bank as published^36^. The schematic at the bottom illustrate the inserting site of TEV recognizing motifs on RAD21. Three repeats of TEV motifs were inserted between P471 and P472 in exon 11 of RAD21. **(b)** TEV cleavage assay using purified nuclei (Upper panel). Briefly, nuclei were purified from HAP1-RAD21^TEV^ cells, plated on poly-Lys plates, and incubated without or with 15U/ml TEV protease at 4C for overnight. Lower panel, western blot analysis of RAD21 from wild-type and RAD21^TEV^ nuclei without and with TEV protease treatment respectively. LMNA was used as the loading control. Numbers indicate the relative amount of intact RAD21. **(c)** Hi-C interaction maps for RAD21^TEV^ nuclei without and with TEV treatment, respectively. Data for the 18-107.3 Mb region of chromosome 14 is shown at 200kb resolution. Bottom, eigenvector E1 across the 18-107.3 Mb region of chromosome 14 at 200kb resolution. **(d)** Saddle plots of Hi-C data binned at 200kb for RAD21^TEV^ nuclei without and with TEV treatment, respectively. The numbers indicate compartment strength. **(e).** Interaction strength of compartments. The bars represent the strength of compartment interactions for each sample as indicated. Dark bars, strength of AA interaction as compared to AB interaction (A-A/A-B). Grey bars, strength of BB interaction as compared to AB interaction (B-B/B-A). **(f)** Hi-C interaction maps for RAD21^TEV^ nuclei without and with TEV treatment, respectively. Data for the 29-34 Mb region of chromosome 14 is shown at 20kb resolution. **(g)** Insulation profiles for the 29-34 Mb regions of chromosome 14 at 20kb resolution. The blue and red lines represent without and with TEV protease treatments, respectively, as in panel C. The panels I, K and K have the same color assignments for the lines as in panel C. The lower panels indicate compartment Eigenvector value E1 across the same region at 200kb resolution. **(h)** Aggregate Hi-C data binned at 20kb resolution at TAD boundaries identified in the sample in NB buffer without TEV treatment. The numbers at the sides of the cross indicate the strength of boundary-anchored stripes using the mean values of interaction frequency within the white dashed boxes., as in Figure 1E. **(i)** *P(s)* plots (upper panels), and the derivatives of *P(s)* plots (lower panels) for Hi-C data from nuclei with or without TEV treatment as indicated. The blue arrows indicate the signature of cohesin loops. (**j**) Aggregated Hi-C data binned at 20kb resolution at loops identified in HAP1 cells (n=8334) by ^14^. Right panel: average Hi-C signals along the blue dashed line shown in the left Hi-C panel. **(k)** Aggregated Hi-C data binned at 20kb resolution at chromatin loops of three different loop sizes, 100-500kb, 500kb-1Mb, and >1Mb. Right panel: average Hi-C signals along the blue dashed line shown in the left Hi-C map in panel J.

We purified nuclei from HAP1-RAD21^TEV^ cells and incubated them overnight with TEV protease at 4°C (Fig. 3b) in a low salt buffer used for nuclei isolation (10mM PIPES, pH7.4, 2mM MgCl2 and 10mM KCl). Western blotting confirmed efficient cleavage of RAD21 (>99% cleaved) as evidenced by the presence of two fragments that could be detected with two antibodies that recognize the N-terminal (Fig. 3b) or C-terminal domain respectively (Extended Data Fig. 3b). Wildtype RAD21 in nuclei purified from parental HAP1 cells was not cleaved.

### RAD21 cleavage reduces CTCF-CTCF loops in isolated nuclei

We performed Hi-C analysis on the HAP1-RAD21^TEV^ nuclei without and with TEV protease treatment in low salt nuclear isolation buffer (NB buffer). Visual inspection of chromosome-wide Hi-C maps did not reveal obvious effects of RAD21 cleavage (Fig. 3c). Compartment domains as detected by E1 were not affected. Compartmentalization strength as quantified by saddle plot analysis (as in Fig. 2c) revealed no change in compartment strength (Fig. 3d-e; Extended Data Fig. 3d-e for a biological replicate). Surprisingly, both TADs and TAD boundary positions, which in cells are dependent on cohesin, showed no changes after cleavage of RAD21 as reflected in near-identical insulation profiles (Fig. 3f-g and see Extended Data Fig. 3f-g for a biological replicate). To quantify insulation strength genome-wide we aggregated interactions at and around TAD boundaries. We find the characteristic pattern of depletion of interactions across the boundaries and line-like features (Fig. 3h). Cleavage of RAD21 did not quantitatively affect these features (Fig. 3h and Extended Data Fig. 3h). We also plotted *P*(*s*) and found no quantitative effects (upper panel, Fig. 3i and Extended Data Fig. 3i). The derivative of *P*(*s*) was also unaffected and indicated the presence of ~100 kb loops even after RAD21 cleavage (arrows in lower panel Fig. 3i and Extended Data Fig. 3i). We conclude that TAD insulation and chromosome folding in general, including formation of many loops is largely intact after RAD21 cleavage.

Finally, to directly assess the effect of RAD21 cleavage on looping interactions between pairs of CTCF sites, we used a list of 8,834 loops previously detected in HAP1 cells ^14^, and aggregated Hi-C data obtained with nuclei in which RAD21 was cleaved and with control nuclei (Fig. 3j and Extended Data Fig. 3j). Visual inspection shows that CTCF-CTCF loops are weakened. We then plotted the Hi-C data along the diagonal line of loop aggregation heatmap as in Fig. 1f. In control nuclei a clear peak is observed that is reduced (~50%) upon RAD21 cleavage. We observed that all loops are somewhat weakened, but that longer-range loops (e.g. >0.5-1 Mb) are less affected (Fig. 3k and Extended Data Fig. 3k).

In summary, cleavage of RAD21 in the low salt buffer used for nuclei isolation leads to reduced looping interactions between CTCF sites, but does not affect maintenance of compartmentalization, boundary insulation, and most loops within TADs that are not positioned at CTCF sites.

### Nuclear retention of cohesin subunits

The limited effect of RAD21 cleavage under low salt conditions suggested that the cohesin complex remains associated with chromatin. We tested this directly by performing a nuclear retention assay followed by Western blotting with antibodies to different cohesin subunits (see Methods). HAP1-RAD21^TEV^ nuclei were plated on poly-Lys plates in low salt NB buffer or NB buffer with 132mM NaCl (NBS1). After overnight incubation in the presence of absence of 15U/ml TEV protease, the supernatant was removed, and nuclear retained proteins were analyzed using Western blotting (see Methods). We find that RAD21, SMC1, SMC3, SA1, SA2, PDS5A, WAPL and NIPBL all are stably associated with nuclei (Fig. 4a, and Extended Data Fig. 4a-c). Upon RAD21 cleavage we did not detect loss of any of these factors from nuclei, and no subunits were detected in the soluble fraction (Figure Extended Data Fig. 4a-c).

**Fig. 4.**
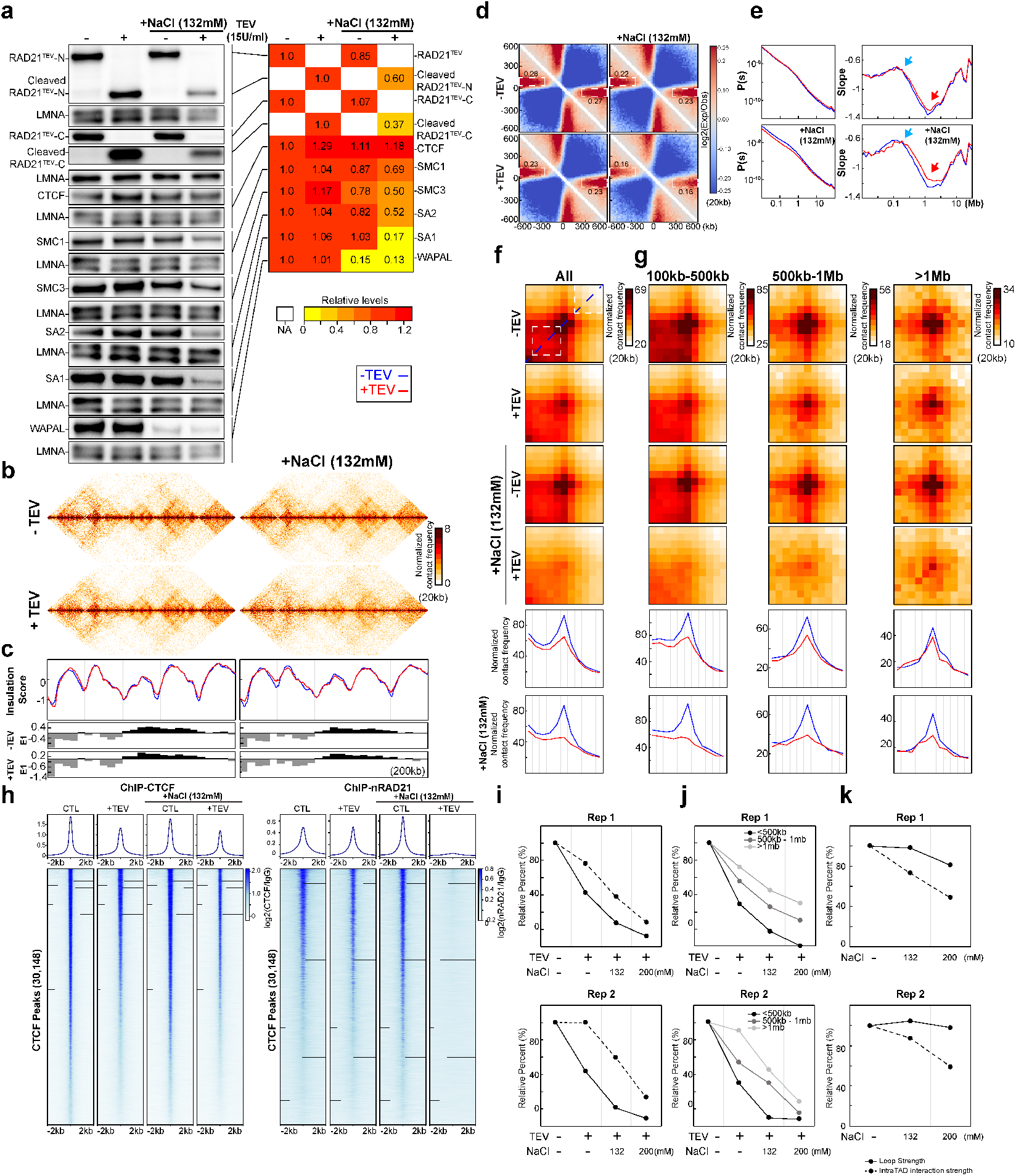
Cleaving RAD21 in NBS1 dissociates cohesin, eliminates CTCF-CTCF loops and reduces intra-TAD interactions. **(a)** Western blot analysis of nuclear retention of cohesin subunits in HAP1-RAD21^TEV^ nuclei with and without TEV protease treatment in NB or NBS1 buffer (see Methods for nuclear retention assay details). LMNA was used as loading controls to normalize each cohesin component in the same lane. Levels of each cohesin subunit in each condition was normalized to its level in nuclei in NB without TEV protease treatment. Relative levels of subunits retained in nuclei are shown in the heatmap on the right. **(b)** Examples of Hi-C interaction maps obtained with HAP1-RAD21^TEV^ nuclei without and with TEV protease treatments in NB (left) or NBS1 (right), respectively. Data for the 29-34 Mb region of chromosome 14 is shown at 20kb resolution. **(c)** Insulation profiles for the 29-34 Mb regions of chromosome 14 at 20kb resolution. The blue and red lines represent without and with TEV protease treatments, respectively. The panels e, f, and g have the same color assignments for the lines as panel c. The lower panels indicate compartment Eigenvector value E1 across the same region at 200kb resolution. **(d)** Aggregate Hi-C data binned at 20kb resolution at TAD boundaries identified in the sample in NB buffer without TEV treatment in NB or NBS1 buffer. The numbers at the sides of the cross indicate the strength of boundary-anchored stripes using the mean values of interaction frequency within the white dashed boxes, as in Fig. 1e. **(e)** *P(s)* plots (left panels), and the derivative of *P(s)* plots (right panels) for Hi-C data from nuclei with or without TEV treatment in NB or NBS1 buffer as indicated. The blue arrows indicate the signature of cohesin loops in each condition. The red arrows indicate the changes in relative contact frequencies at ~2Mb reflecting loop density^32^. **(f)** Aggregate Hi-C data binned at 20kb resolution at loops identified in HAP1 cells (as in Fig. 3j). Lower panels: average Hi-C signals along the blue dashed line shown in the upper Hi-C panels. **(g)** Aggregate Hi-C data binned at 20kb resolution at chromatin loops of three different loop sizes, 100-500kb, 500kb-1Mb, and >1Mb (as in Fig. 3j). Lower panels, average Hi-C signals along the blue dashed line shown in the upper Hi-C map in panel f. **(h)** Profiles of ChIP signals of CTCF and RAD21 at CTCF binding sites without and with TEV treatment in NB or NBS1 buffer. 30148 CTCF binding sites were identified from the CTCF ChIP data of nuclei without TEV treatment in NB. Upper panel, the average CTCF and RAD21 ChIP-seq signals of each condition for 30,148 CTCF binding sites. Lower panel, stack-up heatmap of CTCF and RAD21 ChIP-seq signals for each condition at each of 30,148 CTCF binding sites. **(i)** Quantification of loop strength and intra-TAD interaction strength without and with TEV treatment in NB, NBS1 and NBS2 buffers. Loop strength and intra-TAD interaction strength were calculated as in Fig. 1g. Loop strength and intra-TAD interaction strength from nuclei in NB without TEV treatment were used to normalize respectively. Intra-TAD interaction strength was further normalized by the baseline of 36%, which is observed in the complete absence of cohesin and is the result of general distance decay (see Fig. 1g). Two biological replicates are shown (top and bottom plots). **(j)** Quantification of loop strength for different loop sizes without and with TEV retreatment in NB, NBS1 and NBS2 buffers. Loop strength obtained with nuclei in NB without TEV treatment was used for normalization. Two biological replicates are shown. **(k)** Quantification of loop strength and intra-TAD interaction strength without TEV treatment in NB, NBS1 and NBS2 buffers. Loop strength and intra-TAD interaction strength were normalized as in panel (i).

We next tested cohesin retention under conditions that more resemble physiological ionic strength. We repeated the RAD21 cleavage experiments with nuclei incubated in the NB buffer containing an additional 132mM NaCl (NBS1 buffer). TEV protease cleaved efficiently under these conditions (Fig. 4a, Extended Data Fig. 4a-c). We then performed the same Western blot analysis to assess nuclear retention of cohesin subunits. In the absence of TEV protease we observed some (up to ~20%) reduction in retention of multiple cohesin subunits (RAD21, SMC1, SMC3, SA2, PDS5A), and a larger loss of WAPL (85%). Interestingly, upon RAD21 cleavage we observed an increased loss of nuclear retention of cohesin subunits, and an increase in the soluble fraction (Fig. 4a, Extended Data Fig. 4). However, even under these conditions a sizable fraction of cohesin (~50%) remains stably retained within nuclei. Notably, CTCF is stably retained in low salt and in physiological salt conditions, and no CTCF was detected in the soluble fraction.

### Effect of RAD21 cleavage on chromosome folding under physiological salt conditions

The results in Fig. 4a, combined with the Hi-C analysis in Fig. 3 shows that under low salt conditions, maintenance of chromosome folding does not require intact RAD21 and the cohesin complex remains largely retained within nuclei, and likely remains chromosome associated. However, under physiological salt conditions, cohesin retention and association with chromosomes was affected by RAD21 cleavage. Therefore, we repeated the Hi-C analyses under physiological salt conditions. We incubated HAP1-RAD21^TEV^ nuclei in NBS1 buffer overnight with or without TEV protease and performed Hi-C. E1 was unaffected by RAD21 cleavage. As compared to low salt conditions, compartmentalization was slightly reduced (Extended Data Fig. 5a-c and Extended Data Fig. 6a-c for a biological replicate). Insulation profiles were largely unaffected, but local minima appeared reduced (Fig. 4bc and Extended Data Fig. 6d). To quantify insulation at domain boundaries we again aggregated Hi-C interactions at and around domain boundaries (as above). We observed that insulation at boundaries, and the strength of line-like features, was weakened upon RAD21 cleavage in NBS1 buffer, while in nuclei with intact RAD21 insulation was comparable to that detected under low salt conditions (Fig. 4d and Extended Data Fig. 6e).

Interestingly, *P*(*s*) revealed a decrease in interaction frequency for loci separated by less than 1 Mb (Fig. 4e and Extended Data Fig. 6f). The typical cohesin-dependent shoulder in *P*(*s*) for loci separated by ~100 kb (Figure 1H) was still observed, which was confirmed by plotting the derivative of *P*(*s*): a peak at around ~100 kb was obviously present (Fig. 4e; Extended Data Fig. 6f blue arrows). This indicates that most cohesin-mediated loops are unaffected. We did note that the minimum of the derivative of *P*(*s*) at around 2 Mb is less deep (Fig. 4e, Extended Data Fig. 6f, red arrows), which has been interpreted to reflect a reduced density of loops^32^. Possibly, some loops are lost.

Finally, we quantified the presence of CTCF-CTCF loops (Fig. 4f; Extended Data Fig. 6g for a replicate experiment). In low salt buffer we again noticed that CTCF-CTCF loop strength was reduced but clearly detectable after RAD21 cleavage (Fig. 4f top two heatmaps; this represents an independent biological replicate of the experiments shown in Fig. 3j, and Extended Data Fig. 6g for another replicate). In NBS1 buffer, RAD21 cleavage led to complete loss of CTCF-CTCF loops. We quantified loop strength as described above (Fig. 1f). After RAD21 cleavage, loop strength decreased to ~40% in low salt buffer (as shown in Fig. 3j) and was entirely lost in NBS1 buffer (Figure 4I, quantification for two independent biological replicates). We note that very large loops (>500 kb) could still be detected though their interaction frequency was also greatly reduced (Fig. 4g, 4j). We also calculated the enrichment of intra-TAD interactions as compared to interactions between loci located on either side of the domain. We calculated the average intra-TAD interaction frequency as illustrated in Fig. 1f, and then normalized for the baseline level of 35% detected in the complete absence of cohesin-mediated loops and TADs. This baseline level is due to the expected general distance-dependent decay in interaction frequency (Fig. 1f, 1g). Intriguingly, we found that after RAD21 cleavage, enrichment of intra-TAD interactions was slightly reduced in low salt buffer, and only reduced to ~40-60% in NBS1 buffer (Fig. 4i, two biological replicates). We conclude that RAD21 cleavage under physiological salt conditions leads to complete loss of CTCF-CTCF loops, while enriched interactions within TADs are lost to a much smaller degree. The remaining enriched intra-TAD interactions are likely mediated by the ~50% of cohesin complexes that are still associated with chromosomes under these conditions: first, complete loss of cohesin leads to complete loss of enriched intra-TAD interactions (Fig. 1f, 1g) suggesting the remaining enriched intra-TAD interactions in nuclei with cleaved RAD21 are cohesin-dependent; and second, the *P*(*s*) plot and its derivative indicate that many cohesin-dependent loops are still present after RAD21 cleavage in NBS1 buffer (Fig. 4e, Extended Data Fig. 6f).

Combined, our results strongly predict that under physiological salt conditions, cleavage of RAD21 results in specific loss of cohesin associated with CTCF sites, while cohesins at other locations within TADs remain chromatin-associated and continue to maintain loops. To analyze cohesin association at CTCF sites directly, we performed chromatin immunoprecipitation using antibodies against CTCF and RAD21 using HAP1-RAD21^TEV^ nuclei under different salt conditions and with or without TEV-mediated RAD21 cleavage (Fig. 4h; Extended Data Fig. 7a for replicates). We find that CTCF binding to CTCF sites is not affected by RAD21 cleavage at low salt and at physiological salt concentrations (NBS1 buffer). In contrast, cleavage of RAD21 in NBS1 buffer results in near complete loss of RAD21 ChIP signal at CTCF-bound sites (Fig. 4h, right panel, Extended Data Fig. 7a middle and right panels). At low salt, RAD21 cleavage did not result in loss of RAD21 association at CTCF-bound sites. These observations are consistent with the Hi-C data: RAD21 cleavage only results in complete loss of CTCF-CTCF loops under physiological salt conditions, but these loops remain relatively stable when RAD21 is cleaved under low salt conditions.

Cohesin also associated with active promoters (Extended Data Figure 7B, C, two replicates). Interestingly, when we examined RAD21 binding to active promoters that do not bind CTCF we find that upon cleavage RAD21 is no longer enriched at promoters even at low salt concentrations (NB buffer). This indicates that association of RAD21 with promoters is even more sensitive to RAD21 cleavage than its association with CTCF sites.

### Elevated intra-TAD interactions are sensitive to high salt concentration

Cleaving RAD21 in NBS1 buffer eliminated CTCF-CTCF looping interactions while cohesin-dependent elevated intra-TAD interactions were reduced only 40-60% (Fig. 4i). Given that 50% of SMC1, SMC3 and RAD21 remain on chromosomes, these (cleaved) cohesin complexes may be able to continue to maintain the elevated intra-TAD interactions. We set out to determine whether these remaining cohesin components would dissociate at higher salt concentrations, with concomitant loss of elevated intra-TAD interactions. We repeated the RAD21 cleavage experiments in NB buffer containing 200 mM NaCl (NBS2 buffer). Using the same nuclear retention assay shown in Fig. 4a, we found that even without RAD21 cleavage, up to 50% of RAD21, SMC1 and SMC3 could be dissociated (Extended Data Fig. 8ab, the third lane). After RAD21 was cleaved, the majority of cohesin subunits were no longer retained in the nucleus: more than 70% of RAD21 N-terminus, 90% of RAD21 C-terminus, more than 80% of SMC1, NPBL, SA1 and SA2, and more than 60% of SMC3 were dissociated (Extended Data Fig. 8a). Western blot analysis of released cohesin components in the supernatant confirmed these losses (Extended Data Fig. 8, right panel). CTCF remained bound and was undetectable in the supernatant indicating that CTCF remains stably chromosome-associated in NBS2 buffer.

Next, we performed Hi-C on nuclei incubated in NBS2 in the absence or presence of TEV protease. Incubation of nuclei in NBS2 buffer, in the absence of TEV protease, did not alter the compartment profile (Extended Data Fig. 9a-b and 10a-b; two biological replicates), but compartmentalization strength was slightly reduced (Extended Data Fig. 9bc and 10bc). Insulation profiles also did not change (Extended Data Fig. 9d and 10d), but insulation strength at boundaries was slightly reduced (Extended Data Fig. 9e and 10e). *P*(*s*) showed a decrease in interaction frequency for loci separated by up to ~1Mb (Extended Data Fig. 9f and 10f). Analysis of the derivative of *P*(*s*) showed evidence that in NBS2 buffer there was loss of loops (as seen by the reduced local minimum at around 1 Mb, Extended Data Fig.9f and 10f, red arrows; see above and^32^). Cleavage of RAD21 led to a further weakening of insulation at domain boundaries (Extended Data Fig. 9e and 10e), and loss of more loops as inferred from the derivative of *P*(*s*).

We then investigated CTCF-CTCF loops specifically (Extended Data Fig. 9g and 10g). In the presence of 200 mM NaCl, and with RAD21 intact, CTCF-CTCF loops were slightly reduced (up to 20% in different replicates, Fig. 4k). As expected, cleavage of RAD21 completely eliminated the remaining CTCF-CTCF loops in NBS2, whereas, at low salt concentration loops were still detected (but somewhat reduced, as shown in Fig. 3j and 4f for independent replicates). Different sizes of loops showed similar loss after RAD21 cleavage in NBS2 (Extended Data Fig. 9h and 10h). Quantification of CTCF-CTCF loop strength under the various buffer conditions, and in the absence of TEV protease is shown in Fig. 4k, for two independent replicate experiments: In the presence of intact RAD21, CTCF-CTCF loop strength shows only minor sensitivity to increased salt concentrations up to 200 mM NaCl, while upon cleavage of RAD21 in NBS1 or NBS2 buffer, these loops completely disappear (Fig. 4ij).

Finally, we calculated the enrichment of intra-TAD interactions, as described above (Fig. 4i). In NBS2 buffer, and in the absence of TEV protease, enrichment of intra-TAD interactions was reduced by ~50% (Fig. 4k). Thus, while CTCF-CTCF loops show only very minor sensitivity to salt concentration, enrichment of intra-TAD interactions was much more sensitive. After RAD21 cleavage enrichment of intra-TAD interactions was strongly reduced. As shown in Figure S8, the majority of cohesin was lost from chromosomes under these conditions. The derivative of *P*(*s*) suggests many, but not all loops are lost Extended Data Fig. 9f and 10f). The small number of cohesin complexes (10-30%) that are still retained may mediate these.

We conclude that RAD21 cleavage and increased salt concentration both contribute to loss of cohesin complexes from chromatin, and that these two parameters contribute differently to the loss of the distinct classes of loops: Loops between CTCF sites are highly sensitive to RAD21 cleavage but these interactions show only limited salt sensitivity when RAD21 is intact. In contrast, cohesin-dependent enrichment of intra-TAD loops are sensitive to high salt concentration, and these interactions are much less dependent on intact RAD21.

### Effect of RAD21 cleavage on CTCF-CTCF loops are not due to inter-sister interactions

In the experiments described above, nuclei were purified from non-synchronized cell cultures, and therefore included nuclei in the G1, S and G2 cell cycle stage. During S and G2 cohesin mediates interactions between sister chromatids. Recent analysis of inter-sister interactions using scsHi-C has shown that some CTCF-CTCF interactions can be contacts between sister chromatids that are mediated by cohesive cohesin complexes^37^. Previous studies had shown that RAD21 cleavage can result in loss of inter-sister interactions^8–11^. To rule out that any of our results on the effects of RAD21 cleavage are due to loss of inter-sister instead of intra-sister interactions, we repeated experiments where we sorted G1 nuclei after TEV treatment and formaldehyde fixation of HAP1-RAD21^TEV^ nuclei (Extended Data Fig. 11 and 12, two biological replicates). The results show high concordance with those obtained with unsorted nuclei. Hi-C analysis showed that compartment profiles and compartmentalization strength were only minimally affected by RAD21 cleavage (Extended Data Fig. 11a-c and 12a-c, compare to Extended Data Fig. 5a-c and 6a-c). Insulation at domain boundaries was reduced upon RAD21 cleavage in NBS1 (Extended Data Fig. 11d-e and 12d-e; compare to Fig. 4d and 6d). Analysis of *P*(*s*) and its derivative showed loss of some loops after RAD21 cleavage in NBS1 buffer (Extended Data Fig. 11f and 12f, compare Fig. 4e and Extended Data Fig. 6e). Finally, after RAD21 cleavage, CTCF-CTCF loops were ~60% reduced at low salt conditions, and mostly eliminated at physiological salt concentrations. Enriched intra-TAD interactions were ~60% reduced in NBS1 (Extended Data Fig. 11g-h, 12g-h and 13). When RAD21 was not cleaved, intra-TAD interactions were more salt sensitive than CTCF-CTCF loops. These results confirm all observations made with nuclei isolated from non-synchronized cell cultures and demonstrate that inter-sister interactions were not a major confounding factor.

**Fig. 5.**
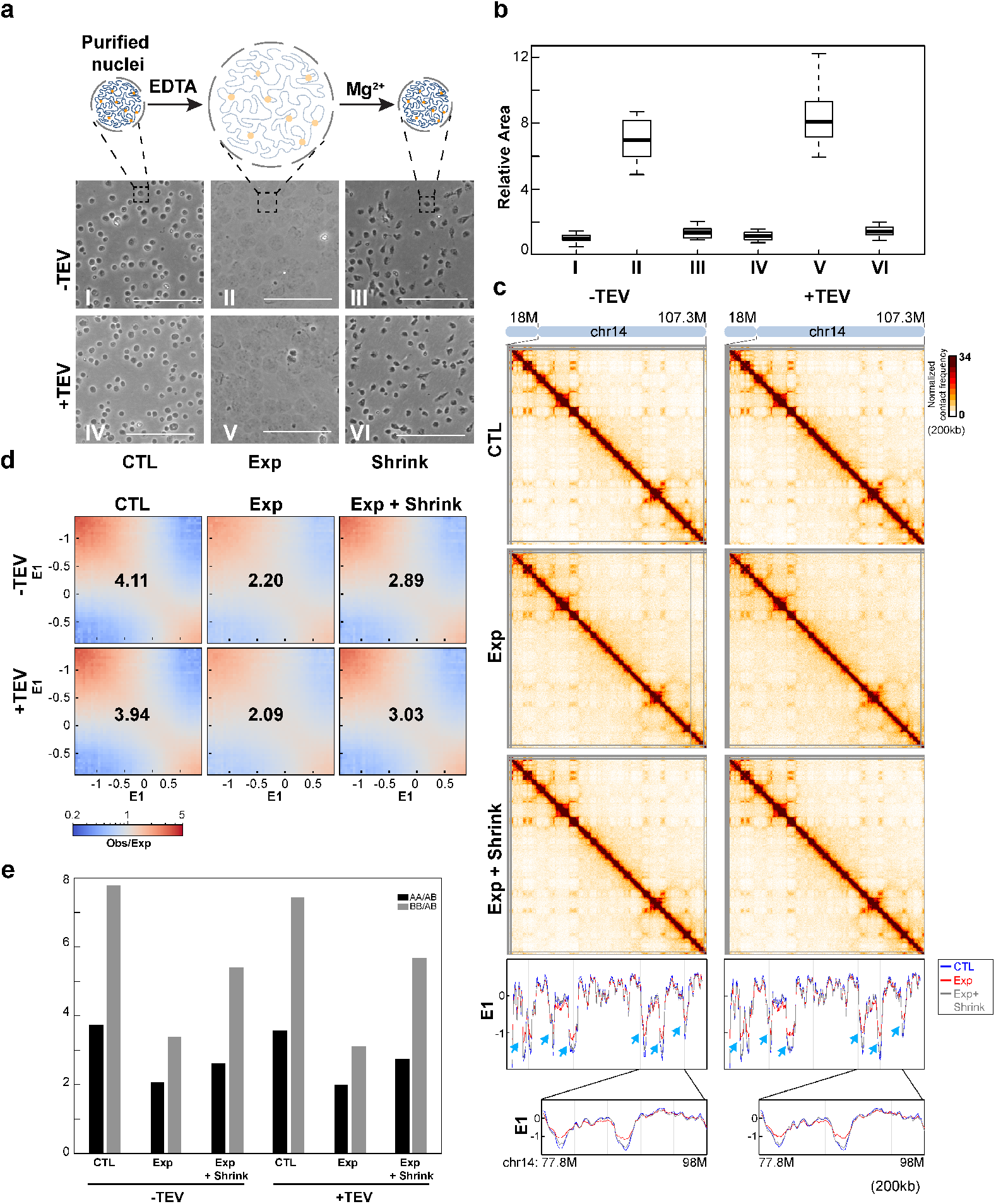
Compartmentalization remains after nucleus expansion and contraction. **(a)**. Changes of nucleus morphology upon expansion and contraction of chromatin (see Methods). Upper, a schematic to show how nuclear morphology changes during chromatin expansion and contraction. Middle and bottom, expansion and contraction for HAP1-RAD21^TEV^ nuclei without and with TEV protease treatments, respectively (Scalebar =100μm). **(b)** Nuclear cross-sectional area changes during the expansion and contraction. The area of 30 nuclei in each condition was measured and plotted using image J and R from the pictures in (a). Median cross-sectional area was set at 1, and fold change in area is shown for expanded and contracted nuclei. **(c)** Hi-C interaction maps for HAP1-RAD21^TEV^ nuclei before expansion, after expansion, and after expansion followed contraction without and with TEV protease treatments, respectively. Data for the 18-107.3 Mb regions of chromosome 14 is shown at 200kb resolution. Bottom panels: Eigenvector E1 across the 18-107.3 Mb regions of chromosome 14 at 200kb resolution. The blue, red and grey lines represent control, expansion and contraction treatments, respectively. Blue arrows indicate E1 changes. Bottom, E1 across the 77.8-98Mb regions of chromosome 14 at 200kb resolution. **(d)** Saddle plots of Hi-C data binned at 200kb for HAP1-RAD21^TEV^ nuclei before expansion, after expansion, and after expansion followed contraction without and with TEV treatments, respectively. The numbers indicate compartment strength. **(e)** Interaction strength of compartments. The bars represent the strength of compartment interactions for each sample as indicated. Dark bars, strength of AA interaction as compared to AB interaction (A-A/A-B). Grey bars, strength of BB interaction as compared to AB interaction (B-B/B-A).

**Fig. 6.**
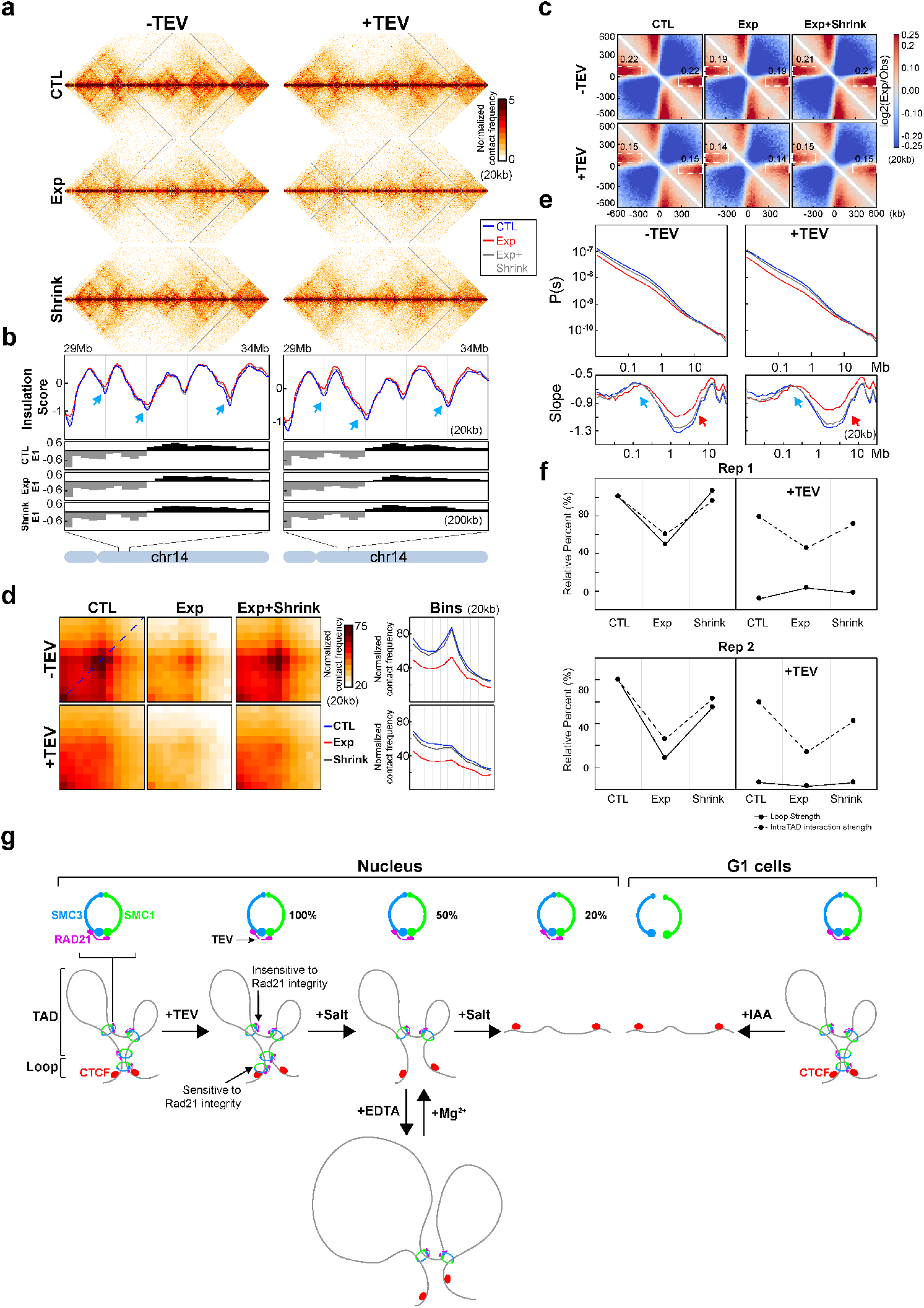
Intra-TAD interactions remain after nuclear expansion and contraction. **(a)** Hi-C interaction maps for HAP1-RAD21^TEV^ nuclei before expansion, after expansion, and after expansion followed contraction without and with TEV protease treatments, respectively. Data for the 29-34 Mb regions of chromosome 14 is shown at 20kb resolution. **(b)** Insulation profiles for the 29-34 Mb region of chromosome 14 at 20kb resolution. The blue, red and grey lines represent control, expansion, and expansion followed by compaction treatments, respectively. The blue arrows indicate weakened insulation boundaries as nuclei were expanded. The lower panels indicate compartment Eigenvector value E1 across the same region at 200kb resolution. **(c)** Aggregate Hi-C data binned at 20kb resolution at TAD boundaries identified in control sample (before expansion) without TEV treatment. The numbers at the sides of the cross indicate the strength of boundary-anchored stripes using the mean values of interaction frequency within the white dashed boxes, as in Fig. 1e. (**d**) Aggregated Hi-C data binned at 20kb resolution at loops as used in Fig. 3j. Right panels: average Hi-C signals along the blue dashed line shown in the left upper Hi-C panel. **(e)** *P(s)* plots (upper panels), and the derivatives of *P(s)* plots (lower panels) for Hi-C data from nuclei before expansion, after expansion, and after expansion followed by compaction with or without TEV treatment as indicated. The blue arrows indicate the signature of cohesin loops in each condition. The red arrows indicate the changes of contact frequency at 2Mb. The reduced minimum in P(s) at ~2 Mb may be due to general loss of interaction for loci up to several Mb, as seen in *P*(*s*), and not due to loss of cohesin loops. **(f)** Quantification of loop strength and intra-TAD interaction strength before expansion, after expansion, and after expansion followed by contraction without and with TEV treatment. Loop strength and intra-TAD interaction strength were calculated as in Fig. 1g. Loop strength and intra-TAD interaction strength from nuclei in NB buffer without TEV treatment was used to normalize. However, distance decay (36%) was not used to normalize intra-TAD interaction strength since distance decay of expanded nuclei in the complete absence of cohesin cannot be calculated here. Two biological replicates were shown. (**g**) Illustration of two biochemically and possibly structurally distance cohesin complexes at positioned CTCF-CTCF loops (dependent on RAD21 integrity) and at loops within TADs (not dependent on RAD21 integrity). The left part with the subtitle of “Nucleus” illustrates HAP1-RAD21^TEV^ in nucleus cleavage experiments, while the right part with the subtitle of “G1 cells” illustrates RAD21 degron experiments using HCT116-RAD21-mAC cells at G1 phase. The percent numbers next to cohesin ring illustrations on top indicate the relative levels of nuclear retained cohesin complex after TEV or salt treatment as indicated in Fig. 4a and Extended Data Fig. 8b.

### Intra-TAD interactions are stable upon chromatin expansion regardless of RAD21 integrity

Our data show that under physiological salt conditions, cohesin-dependent enriched intra-TAD interactions are maintained upon RAD21 cleavage. We wondered whether RAD21 cleavage, and concomitant cohesin ring opening, would render intra-TAD interactions sensitive to mechanical disruption. We recently developed an experimental approach in which chromatin within nuclei can be made to swell extensively by incubating nuclei in a buffer containing 1 mM EDTA. Expansion is caused by the fact that chromatin fibers become decondensed in the presence of EDTA and their contour length (nm/kb) increases^38, 39^. Nuclei can then be shrunk to their original size by replacing the buffer with HBSS which will cause the chromatin fiber to condense. We reported that chromosome folding is highly elastic during expansion and contraction cycles^39^.

Expansion and contraction involve chromatin movement, as expansion is prevented by fixation, and therefore physical forces may be exerted on looping interactions mediated by cohesin. We used this approach to determine whether dramatic expansion of chromatin would irreversibly disrupt cohesin-dependent enriched intra-TAD interactions when RAD21 is cleaved. We plated isolated HAP1-RAD21^TEV^ nuclei on poly-D-lysine coated plates in HBSS buffer at 4°C (Fig. 5a, Extended Data Fig. 15a for a biological replicate). When we replaced the buffer with expansion Buffer (EB, 1mM EDTA, 10mM HEPES, pH 7.4) we observed that nuclei expanded extensively. Microscopic analysis showed that the cross-sectional area of nuclei increased around ~7-fold, representing a ~18-fold increase in nuclear volume (Fig. 5b). When we then replaced the buffer with HBSS, nuclei contracted to very close to their original size (1-1.2x their original size). We repeated these experiments after overnight incubation with TEV and found that expansion and contraction of nuclei was unaffected.

We first set out to ensure that all results on the effects of RAD21 cleavage described above in NBS1 buffer were reproduced when nuclei were incubated in the HBSS buffer used for the expansion – contraction experiments. HBSS and NBS1 are comparable in terms of their ionic strength. We again performed nuclear retention assays to determine whether cohesin subunits became solubilized after cleavage (Extended Data Fig. 14a). We find that after RAD21 cleavage in HBSS buffer, about 50% of cohesin subunits were lost from nuclei, which is comparable to what we observed when nuclei were incubated in NBS1 buffer (compare with data shown in Fig. 4a). We then performed Hi-C on HAP1-RAD21^TEV^ nuclei in HBSS buffer in the presence or absence of TEV protease (Extended Data Fig. 14 for two replicates). Compartmentalization strength was not affected by RAD21 cleavage (Extended Data Fig. 14b-d). Insulation at domain boundaries was somewhat reduced (Extended Data Fig. 14ef). *P*(*s*) was largely unchanged (Extended Data Fig. 14g), but CTCF-CTCF loops were strongly reduced (Extended Data Fig. 14h). We conclude that all results obtained with NBS1 buffer are reproduced in HBSS buffer.

We then performed Hi-C on isolated nuclei without expansion (4 hours incubation in HBSS), on expanded nuclei (4 hours in EB, 1mM EDTA, 10mM HEPES, pH 7.4), and on nuclei that we were first expanded for 4 hours, and then contracted for 1 hour again in HBSS (Fig. 5, see Extended Data Fig. 15 for a biological replicate). In these experiments RAD21 is intact. E1 profiles did not change, but compartmentalization strength was reduced when nuclei were expanded. The B-compartment was most affected (Fig. 5cde and Extended Data Fig. 15bcd). When nuclei were contracted again, compartmentalization became stronger again but did not reach the full strength observed before expansion. Insulation profiles were mostly unchanged, but we did notice a slight loss of insulation at domain boundaries (Fig. 6a left panel, b left panel, blue arrows; Fig. 6c top panel; Extended Data Fig. 16 left panels). CTCF-CTCF loops, and enriched intra-TAD interactions became weaker upon expansion, and both regained full strength after nuclei were contracted again (Fig. 6d upper panels). This weakening of interactions in expanded nuclei is likely the result of a general decrease in interaction frequency of loci separated by up to 1 Mb, as readily observable in *P*(*s*) plots (Fig. 6e left panels, Extended Data Fig. 16c left panels). This general decrease in interaction frequencies can be explained by the fact that the contour length of the chromatin fiber (nm/kb) increases in expansion buffer^38–40^. Given that CTCF-CTCF looping interactions as well as enriched intra-TAD interactions regain full strength after contraction of nuclei, it is likely that these interactions were never lost upon expansion.

We performed the same analyses on nuclei in which RAD21 was cleaved before nuclei were expanded and contracted again. As described above for NBS1 buffer, after RAD21 cleavage in HBSS buffer before expansion insulation at domain boundaries is reduced and CTCF-CTCF loops were mostly lost, while enriched intra-TAD interactions were still detected (Fig. 6d lower panel and Extended Data Fig. 16d lower panel). In expanded nuclei enriched intra-TAD interactions were reduced, but these were restored when nuclei were contracted again (Fig. 6d lower panel, Extended Data Fig. 16d lower panel, and 6F right panel). Derivative of *P*(*s*) plots confirm that during expansion and contraction of nuclei the signature of cohesin-dependent loops remains present (Fig. 6e, Extended Data Fig. 16c bottom panels, arrows). We conclude that intra-TAD interactions are maintained upon extensive nuclear expansion and contraction even when RAD21 is cleaved.

## DISCUSSION

Our results show that cohesin complexes can associate with chromatin and mediate looping interactions in different ways dependent on their genomic location (Figure 6g). At CTCF sites, stable loops require closed cohesin rings that may (pseudo-)topologically embrace DNA. Within TADs, cohesin complexes can maintain loops even when the ring is opened by RAD21 cleavage. The results point to structural alterations of the complex, and possibly subunit exchanges, as cohesin loads on chromatin, travels along chromatin to extrude loops, and encounters CTCF-bound sites.

The observation that the cohesin complex forms rings has led to a model that the complex can mediate sister chromatid cohesion by entrapping the two sister DNA molecules within the ring ^6, 7^. Biochemical studies where the subunits of the complex that make up the ring can be covalently crosslinked after the complex has established cohesion between two circular DNA molecules, have shown that cohesion is then resistant to protein denaturation, providing strong evidence that the two DNA molecules are topologically trapped within such artificially covalently closed cohesin rings^6, 7^. This model explains the observation that proteolytic cleavage of the RAD21 subunit is sufficient to release sister chromatids at the metaphase-to-anaphase transition^8–11^. The model that cohesin forms a ring around DNA is also consistent with the observation that RNA polymerase can push the complex towards the end of genes.^41–43^

In recent years it has become evident that besides its role in sister chromatid cohesion, the cohesin complex has additional roles in chromosome folding. The cohesin complex can extrude chromatin loops all along chromosomes. While many loops are dynamically formed and not positioned at specific reproducible genomic positions, blocking of extrusion at CTCF sites results in a subset of loops that are specifically anchored at CTCF sites. How cohesin mediates loop formation, and whether it involves similar modes of association with DNA as occurs when cohesin mediates cohesion is not known. Recent single molecule experiments using covalently closed cohesin rings suggest that loop extrusion can occur without the need for ring opening at any point during the loading and extrusion process, pointing to potential differences in cohesin-DNA interactions during cohesion and extrusion^15^. Several structures of (parts of) the cohesin complex in association with DNA have been solved, and together with loop extrusion simulations these suggest that the cohesin ring may not need to open during the process of loop formation^36, 44, 45^.

Chromatin loops are readily detected by Hi-C in two ways: first, dots of enriched interactions in Hi-C contact maps represent positioned loops, often at CTCF sites. Second, *P*(*s*) plots of Hi-C interaction frequencies shows a characteristic shoulder at ~100 kb that represent loops throughout the genome, that are not positioned at reproducible genomic locations and are mostly contained somewhere between CTCF sites leading to TAD formation. Both these Hi-C features completely disappear when RAD21 is efficiently destroyed using degron approaches^23, 25^. We show that cohesin ring opening through precise RAD21 cleavage results in loss of positioned loops at CTCF sites, while other cohesin-dependent loops within TADs are maintained. When RAD21 was cleaved in low salt buffer, CTCF-CTCF loops were reduced by about 50%. Chromatin binding assays showed minimal loss of cohesin subunits from chromatin. Interestingly, ChIP showed that after cleavage, RAD21 binding to CTCF sites was not reduced under low salt conditions. This indicates that CTCF-CTCF looping interactions are more sensitive to RAD21 cleavage than RAD21 chromatin binding at CTCF sites. At promoters, cleavage of RAD21 led to loss of RAD21 binding even at low salt concentrations. Possibly the interaction between cohesin and CTCF is stable enough under low salt conditions to maintain cohesin at CTCF sites even after RAD21 is cleaved. Under physiological salt conditions, cleavage of RAD21 results in complete loss of CTCF-CTCF loops, loss of RAD21 binding at CTCF sites and ~50% decrease in chromatin bound cohesin complexes. Previously we showed that fragmenting chromatin with a restriction enzyme results in loss of CTCF-CTCF loops and loss of ~50% of chromatin-bound cohesin complexes^35^. Therefore, CTCF-CTCF loops and cohesin association with CTCF sites are abolished when either the DNA is fragmented, or the cohesin complex is cleaved. Taken together these observations suggest that at CTCF sites, and possibly promoters, DNA is passing through the cohesin ring. Further, CTCF-CTCF loops and cohesion are both sensitive to RAD21 cleavage, pointing to similarities in the way cohesin associates with DNA as it stabilizes loops at CTCF sites, and when it mediates cohesion between sister chromatids.

Cohesin-dependent interactions within TADs were stably maintained after RAD21 cleavage but, in contrast to CTCF-CTCF loops were salt sensitive even when RAD21 was intact. This suggests that when cohesin mediates loops within TADs, e.g., when it is actively extruding while not blocked at CTCF sites, it is associated with DNA in a different way, at least during some steps of the extrusion process, as compared to when it mediates loops between CTCF sites. The salt sensitivity of DNA binding of this subpopulation of cohesin complexes suggests they do not encircle DNA. A cryo-EM structure of the cohesin complex in the presence of NIPBL, which is required for extrusion ^15^, shows that DNA is tightly held between NIPBL, RAD21 and the SMC head domains ^44^. Modeling suggests cohesin’s tight grip on DNA is part of a cycle of dynamic changes in cohesin that drives loop extrusion^44, 45^. A tight binding to DNA can explain why cleaved cohesin can still stably maintain loops within TADs even after dramatic nuclear expansion and contraction. Clearly, cohesin associates with DNA, and mediates loops in distinct ways when it is located at CTCF sites or when it is located within TADs. Our results that dynamically extruding cohesin complexes within TADs maintain loops after RAD21 cleavage and possibly do not entrap DNA are consistent with recent finding in *S. cerevisiae*: Srinivasan and co-workers showed that a mutant cohesin complex (smc1DDsmc3AAA) that cannot entrap DNA can still load onto and move along chromatin^7^. Our interpretation that cohesin can associate with DNA in two ways, one dependent on ring integrity (at CTCF-CTCF loops) and one that does not (within TADs), is consistent with Srinivasan’s conclusion that cohesin in yeast can associate topologically (to mediate cohesion) and non-topologically (to associate and move along chromatin). Finally, our data unify and explain apparently disparate results in the literature where complete RAD21 degradation or RAD21 cleavage have quantitatively different effects on chromatin looping ^25, 27^.

Assuming that cohesin that mediates loops within TADs is actively extruding until it encounters CTCF sites, our data suggest that extruding cohesin initially holds DNA tightly but then must alter its conformation and the manner in which it associates with DNA upon engaging with CTCF. Possibly this involves subunit exchange and/or ring opening and closing to establish cohesin complexes that hold pairs of CTCF sites together in a manner that is related to how cohesin holds pairs of sister chromatids. Future studies can address this aspect of cohesin dynamics.

The semi-in vitro approach we employed here using purified nuclei that contain modified cohesin complexes that can be precisely controlled in combination with Hi-C analysis may provide a way to study loop extrusion and the dynamic changes in the cohesin complex in a highly controlled fashion. For instance, it may be possible that by adding or omitting nucleotides or ATP active processes such as transcription and loop extrusion may be experimentally controlled and effects of chromatin looping can then be studied. An advantage of such approach over more conventional degron approaches, is that confounding factors such as cell cycle progression or arrest during the experiments, which we found can obscure direct effects of cohesin depletion, can be minimized.

## Materials and method

### Cell culture

HCT-116-RAD21-mAID-mClover cells were kindly provided by Natsume et al., 2016 ^46^. These cells were cultured in McCoy’s 5A medium, GlutaMAX supplement (Gibco, 36600021) supplemented with 10% FBS (Gibco, 16000044) and 1%penicillinstreptomycin (Gibco, 15140) at 37°C in 5% CO2. HAP1 cells were purchased from Horizon Genomics (Cambridge, UK). Both HAP1 and HAP1^RAD21TEV^ cells were cultured in IMDM medium, GlutaMAX supplement (Gibco, 31980097) supplemented with 10% FBS (Gibco, 16000044) and 1% penicillin-streptomycin (Gibco, 15140) at 37°C in 5% CO_2_.

### Cell cycle analysis and G1 cell sorting

For cell cycle analysis, cells were washed using 1xPBS once and fixed in 90% ethanol at − 20°C for at least 24 hours. Fixed cells were washed in 1xPBS and then resuspended in PBS containing 2mM MgCl_2_, 0.5mg/ml RNase A(Roche, 10109169001), 0.2uM FxCycle far red (200uM stock in DMSO, Thermo Fisher F10348). The samples were incubated at 20°C for 30min and then analyzed using an LSRII flow cytometry instrument with red and green channels to monitor DNA contents and GFP signals for RAD21 levels, respectively.

To arrest cells in G1 phase, we first treated the cells with 2mM thymidine for 12 hours to arrest cells in S phase, then fresh medium was added after washing the cells with PBS twice. After growing in fresh medium for 12 hours, the cells were treated with 400uM mimosine for 12 hours, and the cells were arrested at the boundary of G1/S. To sort G1 cells for Hi-C analysis, cells were fixed following the Hi-C protocol using 1% FA. Fixed cells were washed in 1xPBS and then resuspended in PBS containing 2mM MgCl_2_, 0.1% Saponin, 0.5mg/ml RNase A(Roche, 10109169001), 0.2uM FxCycle far red (200uM stock in DMSO, Thermo Fisher #F10348). The samples were incubated at 20°C for 30min then analyzed using an BD FACS Aria II flow cytometry instrument with red and green channels to monitor DNA content and RAD21 levels, respectively. To avoid obtaining any cells in S phase, only the cells in left part of G1 peak were collected (red dashed box in Supplementary Figure 3). FACS data were processed and analyzed using FlowJo v.3. Viability gates using forward and side scatter were set on each sample. DNA content was plotted as a histogram of the red channel while RAD21 level was plotted as a histogram of the green channel.

### Creation of HAP1-RAD21^TEV^ cells

The pSpCas9(BB)-2A-Puro (PX459) v.2.0 plasmid (F. Zhang lab) was obtained from Addgene (#62988) and used to construct the CRISPR/Cas vector according to the protocol of Ran et al. ^47^ The sequence of the gRNA used to generate HAP1-RAD21^TEV^ is CTCATCTATGTTTGTTCTGC.

To construct a donor plasmid for insertion of tandem TEV motifs, a pCMV-based plasmid was constructed using the following three templates:

- As below, the sequence including three TEV motifs was inserted between chr8: 117,864,243 and 117, 864, 244. This sequence was obtained from ^48^ AGGGCTAGAGAGAATTTGTATTTTCAGGGTGCTTCTGAAAACCTTTACTTCCAAGGAGAGCTCGAAAATCTTTATTTCCAGGGAGCTAGC
- 5’ homology arm of insert site (2,027bp, chr8:117,862, 217 – chr8:117,864,243),
- 3’ homology arm of insert site (2,032bp, chr8: 117,864,244 – chr8: 117, 866, 275)

The genomic co-ordinates are from hg19, and T at chr8:117,864,244 was mutated to A to create the AvrII digestion site for cloning and genotyping purpose. This changes codon CCA (Proline) to CCT (proline).

Genomic DNA from HFF1 cells was used as template for homology arms, which were amplified using Q5 High-Fidelity DNA Polymerase (New England Biolabs, #M0491). Both homology arms and TEV motif sequence were cloned into pCMV plasmid by Gibson assembly using NEBuilder HiFi DNA Assembly Master Mix (NEB, #E2621) and the sequence was confirmed by Sanger sequencing.

To generate stable cell lines, 0.5 × 10^6^ cells were plated and transfected with CAS9-gRNA and linearized donor plasmids using TurboFectin 8.0 (OriGene, USA), following the instructions. Before transfection the medium was changed with antibiotics-free medium containing 0.1uM SCR7 ^49^. Twenty four hours after transfection, 2 μg/ml of puromycin was added and, 2 days later, the surviving cells were diluted in 96-well plates to obtain single cell derived colonies for further screening. HAP1-RAD21^TEV^ cells genotyped by PCR and were further confirmed using a TEV cleavage assay (see below).

### Nucleus purification and TEV cleavage

Nuclei were purified according to Sanders et al ^39^. Briefly, HAP1-RAD21^TEV^ cells were trypsinized, collected in medium, and counted. Around 100million cells were collected and washed in cold PBS twice, and once in NB buffer (10mM PIPES pH 7.4, 2mM MgCl_2_, 10mM KCl, protease inhibitor (PI, ThermoFisher, #78438)). The cells were then resuspended in NB buffer with 0.1% NP-40, 1mM DTT, and protease inhibitors, and incubated on ice for 10mins. The cells were lysed using a Dounce homogenizer with pestle A and then loaded onto a sucrose cushion (NB buffer + 30% sucrose + 1mM DTT and 5ml of cell lysis for 20ml sucrose cushion), and finally centrifuged at 800g for 10mins. The nucleus pellets were washed once in cold NB buffer, then resuspended in NB and counted. Around 4 or 10 million nuclei were plated onto 60mm or 100mm poly-D-lysine culture dishes, respectively (Corning BioCoat, #356469 and #354468), and incubated at 4°C overnight. For the TEV cleavage assay, 150 units of TEV enzyme (AcTEV Protease, Thermo Scientific, 12575-015) were added to 10 million nuclei in 10ml buffer (15U/ml TEV) before the nuclei were plated onto the dishes and kept at 4°C overnight. Control plates contained no TEV, otherwise TEV concentrations were as indicated in Figures.

### Hi-C for sorted G1 nuclei

Nuclei were purified as described above. Around 30 millions of nuclei were resuspended in 14ml NB or NB + 132mM (NBS1) buffers in 15ml falcon tubes. To the samples where RAD21 was to be cleaved, 15U/ml of TEV protease was added. All tubes were gently mixed well and incubated at 4C overnight. After fixation in 1% formaldehyde (FA), nuclei were washed once with PBS and then stained using propidium iodide (PI). G1 nuclei were then sorted using an BD FACS Aria II flow cytometry instrument with red channel to monitor PI (DNA content). To avoid obtaining any S phase cells, only the cells in left part of the G1 peak were collected as described above. Around 1million G1 nuclei were collected and washed twice using 500ul cold NEBuffer 3.1. Cautions should be taken when nuclei were spun down at 1000g for 5minutes. After wash, nuclei were resuspended in 300ul NEBuffer 3.1 with 0.1% SDS and incubated in 65°C for 10min. The following steps are the same as below for DpnII digestion.

### Hi-C for expanded and contracted nuclei

Nucleus expansion experiments were performed as described in^39^. Briefly, after nuclei were attached to poly-D-lysine coated dishes overnight at 4°C, nuclei were washed once with HBSS, then quickly washed twice with Expansion Buffer (EB: 1mM EDTA, 10mM HEPES, pH 7.4). After washing, nuclei were kept in 10ml EB at room temperature for 4 hours before nuclei were fixed in 1% FA for Hi-C analysis. In the control plates, the nuclei were also washed for 4 times using HBSS buffer and kept in HBSS buffer for 4 hours. For nuclei contraction experiments, nuclei were first expanded in EB buffer for 4 hour as described above, then EB buffer was exchanged for HBSS buffer and nuclei kept at room temperature for another 1 hour before fixation with 1% FA for Hi-C analysis.

### Hi-C experiments

Hi-C for fixed cells was performed as described previously^50^. For the nuclei treated as described above, Hi-C protocol was modified. Briefly, for the nuclei attached on the plate, the buffer used in each condition was replaced with the same buffer but containing 1% FA, then the plates were kept at room temperature for 10mins with a few times of gently shaking. After 10 minutes, 2% of 2.5M glycine was added to quench FA for 5 minutes, and the plates were kept on ice for at least 15 minutes. The fixed nuclei were then scratched from the plates and washed twice using NEBuffer 2. For the nuclei using HindIII digestion, from this point, the following steps are the same with that for the cells. Briefly, 35ul of 1% SDS was added to 312ul nuclei in NEBuffer 2 and the tube was gently mixed. The tube was then incubated in 65°C for 10min and then put on ice immediately. Before 400U HindIII was added, 40ul of 10% Triton X-100 was added and gently mixed. HindIII digestion was performed at 37°C overnight with gently rocking. For the nuclei using DpnII digestion, after cold NEBuffer 3.1 washing twice, 300ul NEBuffer 3.1 with 0.1% SDS was added and then the nuclei were scratched off the plate and collected. The nuclei in NEBuffer 3.1 + 0.1%SDS were then directly incubated in 65°C for 10mins and the n put on ice immediately. Before 400U DpnII was added, 40ul of 10% Triton X-100 was added and gently mixed. DpnII digestion was performed in 37°C for overnight with gently rocking. Once enzyme digestion was done, the reaction was incubated at 65°C for 10mins to inactivate HindIII or DpnII. After this, the DNA overhanging ends were filled in with biotin-14-dCTP for HindIII digested chromatin, or biotin-14-dATP for DpnII digested chromatin, at 23 °C for 4 hours and then ligated with T4 DNA ligase at 16 °C for 4 hours. DNA was treated with proteinase K at 65 °C overnight to remove cross-linked proteins. Ligation products were purified, fragmented by sonication to an average size of ~200 bp and size-selected to fragments of 100–350 bp. We then performed end repair and dA-tailing and selectively purified biotin-tagged DNA using streptavidin beads. Illumina TruSeq adapters were added to form the final Hi-C ligation products, samples were amplified and the PCR primers were removed. Hi-C libraries were then sequenced using PE50 bases on an Illumina HiSeq 4000 instrument.

### Hi-C and data analysis

Hi-C PE50 fastq raw sequencing files were mapped onto hg19 human reference genome using distiller-nf mapping pipeline (https://github.com/mirnylab/distiller-nf). After mapping, aligned reads were further processed to remove duplicates (https://github.com/mirnylab/pairtools) to obtain a set of filtered reads defined as valid pairs. Valid pairs were then binned into contact matrices at 20 kb and 200 kb resolutions using cooler50. Intrinsic Hi-C biases were removed using the Iterative Correctionm and Eigenvector decompositiom (ICE) procedure^51^ was applied to all of the matrices, ignoring the first two diagonals to avoid short-range ligation artefacts at a given resolution, and bins with low coverage were removed using the MADmax filter with default parameters. Contact matrices were stored in ‘.cool’ files and used in downstream analyses.

For downstream analyses using cWorld scripts, (https://github.com/dekkerlab/cworld-dekker) cooler files were first converted to matrix files using cooltools dump_cworld (https://github.com/mirnylab/cooltools). The matrix files were then scaled to 1 million reads using cworld scaleMatrix before other analyses. For contact matrix visualization of a region, the matrix of this region was first extracted using cworld extractSubMatrices and the heat map of the region was generated using cworld heatmap.

For aggregation of loop interactions, the previously identified sets of HCT-116-RAD21-mAC and HAP1 looping interactions were used, respectively. In total, 3169 and 8334 looping interactions are on the structurally intact chromosomes of HCT-116-RAD21-mAC and HAP1, respectively. To visualize the looping interactions, we aggregated 20 kb binned data at all loops using cworld interactionPileUp. We also aggregated 20kb binned data at different sizes of loops, 100kb-500kb, 500kb-1Mb and greater than 1Mb. The size of a loop refers to the distance between the two loop anchors.

For *P*(s) plots and derivatives, the cis reads from the valid pairs files were used to calculate the contact frequency *(P)* as a function of genomic separation (s) (cooltools). All of the *P*(*s*) curves were normalized for the total number of valid interactions in each dataset. Corresponding derivative plots were calculated from each *P*(s) plot.

For interaction aggregation at TAD boundaries, we first calculated observed/expected Hi-C matrices of each sample for 20 kb binned data, correcting for average distance decay as observed in the *P*(s) plots (cooltools compute-expected). We then aggregated the observed/expected Hi-C matrices of each sample at the TAD boundaries that were identified from the sample without any treatments, covering 600kb up and downstream of each boundary, and then generated a pileup heatmap of TAD boundaries for each sample.

For compartment analysis, compartment boundaries were identified in cis using eigen vector decomposition on 200 kb binned data with the cooltools call-compartments function. A and B compartment identities were assigned by gene density tracks such that the more gene-dense regions were labelled A compartments, and the PC1 sign was positive. Changes in compartment type therefore occur at locations where the value of PC1 changes sign. Compartment boundaries were defined at these locations, except for when the sign change occurred within 400 kb of another sign change. We noticed that translocation between chr9 and chr22 in HAP1 cells affects compartment assignment on chr9, thus we excluded chr9 for the subsequent compartment analysis for all HAP1 cells.

To measure compartmentalization strength, we calculated observed/expected Hi-C matrices for 200 kb binned data, correcting for average distance decay as observed in the *P*(*s*) plots (cooltools compute-expected). We then arranged observed/expected matrix bins according to their PC1 values of the sample without any treatments in each replicate. We aggregated the ordered matrices for each chromosome within a dataset and then divided the aggregate matrix into 50 bins and plotted, yielding a saddle plot (cooltools compute-saddle). Strength of compartmentalization was defined as the ratio of (A−A + B−B)/(A−B + B−A) interactions. Strength of A-A and B-B interactions were separately calculated using AA/AB and BB/BA, respectively. The values used for this ratio were determined by calculating the mean value of the 10 bins in each corner of the saddle plot.

### Western blot for cohesin components

For each condition, 4 millions of nuclei were plated on 60mm poly-Lysine coated plates in 4ml buffer at 4°C for overnight. To analyze released cohesin proteins, 3ml buffer was collected and spun at 800g for 10 minutes to remove unattached nuclei. The supernatants were then concentrated using Amicon columns (3KDa, #UFC500396). After concentration, the final volume of each concentrated sample was adjusted to 200ul and 50ul of 5× sample buffer (ThermoFisher, #39000) was added. The samples were then boiled at 100°C for 5 minutes before western blot analysis. To analyze cohesin proteins retained in nuclei, all buffers were completely removed from each plate and 200ml RIPA buffer (Thermo Fisher, #89900) containing protease inhibitor and TurboNuclease (Accelagen, #N0103M) were added to each plate. All the plates were then incubated at 4°C for 10minutes and the lysed nuclei were scratched and collected. After spun at 8000g for 10 minutes, the supernatant was transferred to a new tube and 5x sample buffer was added. After mixing, the lysis was boiled at 100°C for 5mins for western blot analysis.

The volume for approximately the same number of cells or nuclei for each sample was loaded into each lane of a protein gel for separation. Two types of protein gel and buffer were used. To separate small proteins (MW<50KD), NuPAGE 4–12% Bis-Tris protein gels (Thermo Fisher, #NP0322BOX) was used with NuPAGE MOPS SDS Running Buffer (Thermo Fisher, #NP0001). For large proteins (MW > 100kD), NuPAGE 3-8% TrisAcetate protein gels (Thermo Fisher, #EA03752BOX) were used in NuPAGE Tris-Acetate SDS running buffer (Thermo Fisher, #LA0041). Proteins were transferred to nitrocellulose membranes (Bio-Rad, #1620112) at 30 V for 2 h in 1× transfer buffer (Thermo Fisher, #35040) in the cold room. The membranes were blocked with 5% milk in TBST (20mM Tris-HCl, pH 7.4, 150mM NaCl and 0.1% Tween-20) for 30 minutes at room temperature. The membranes were then incubated with the specified primary antibodies diluted 1:1,000 in TBST overnight at 4°C. The membranes were washed three times with TBST for 10 min at room temperature each, then incubated with secondary antibodies (anti-rabbit IgG HRP-linked, Cell Signaling, 7074) diluted 1:5,000 in TBST for 2 hours at room temperature. The membranes were then washed three times with PBS-T for 10 min each. Then, the membranes were developed and imaged using SuperSignal West Dura Extended Duration Substrate (Thermo, #34076) and Bio-Rad ChemiDoc.

### Nucleus ChIP-seq experiments

ChIP-seq experiments were based on the protocol in our recent work with some minor modifications^52^. Briefly, for each condition, 40 millions of nuclei were plated on two 100mm poly-Lysine plates. After FA fixed, nuclei were washed twice using washing buffer (20mM TrisHCl pH8.0 + protease inhibitor), then 900ul sonication buffer (20mM Tri-HCl pH 8.0, 0.2% SDS, 0.5% sodium deoxycholate, and protease inhibitor) was added to each plate and the nuclei were scraped and collected. The chromosomes were sonicated to fragments around 200-500bp using BioRaptor Pico (30sec ON, 30sec OFF, 8 cycles). After spinning at 16,000g for 10min, the fragmented chromatin in the supernatant was split into aliquots of 5 millions nuclei per tube and diluted in 1200ul IP buffer with final 0.1% SDS concentration (20mM Tri-HCl pH 8.0, 150mM NaCl, 2mM EDTA, 0.1% SDS, 1% Triton-100, and protease inhibitor). For each tube, around 4ug of antibody was added and for each antibody, 10 million nuclei were used for ChIP. The primary antibodies used in this study included, CTCF (Millipore, #07-729), RAD21 (Abcam, #ab154769, recognizes N-terminal of RAD21), RAD21 (Abcam, #ab992, recognizes C-terminal of RAD21), and Rabbit IgG (Sigma, #I-5006). Chromatin was incubated with primary antibodies on a rocker at 4°C for overnight. After rewashing with IP buffer, 20ul of Dynabeads Protein G (Thermo Fisher, #10004D) were added to each tube followed by incubation on a rocker at 4°C for 2 hours. After the beads were washed for three times, immunoprecipitated DNA was eluted and 5ng ChIP DNA was used to prepare sequencing libraries in the same way as described for Hi-C above.

### ChIPseq data analysis

Sequencing reads were mapped onto hg19 using Bowtie 2. HOMER was used to clean mapped reads, examine quality of ChIP experiments, call peaks and generate visualization files^53^. Both profile and cluster plots of ChIPseq signals were generated using DeepTools^54^. Lists of active TSSs in HAP1 cells were obtained from our recent work^52^. Briefly, publicly available active mark H3K4me3 ChIP-seq signal was used to rank annotated TSSs from the most active to inactive^55, 56^. Top 13, 412 TSS were selected as Active TSSs in HAP1. For stacking analysis in Extended Fig. 7b and 7c, TSSs overlapped with CTCF peaks were used as TSS with CTCF peaks, while TSSs that are at least 2kb away from a CTCF sites were used as TSS with no CTCF peaks.

### Resources and data

All antibodies for Western blot and ChIPseq experiments are listed in Supplemental Table 1. All ChIPseq and Hi-C libraries are listed in Supplemental Table 2 and 3.

## Author contributions

Y.L. and J.D. conceived and designed the project. Y.L. performed all the experiments and analyzed all the data. Y.L. and J.D. wrote the manuscript.

## Acknowledgement

We thank all the members of the Dekker laboratory for discussion, and Rachel McCord for advice on the nuclear expansion assay. We acknowledge support from the National Institutes of Health Common Fund 4D Nucleome Program (DK107980) and the National Human Genome Research Institute (HG003143). J.D. is an investigator of the Howard Hughes Medical Institute.

## REFERENCES

1. Haering, C.H., Lowe, J., Hochwagen, A. & Nasmyth, K. Molecular architecture of SMC proteins and the yeast cohesin complex. Mol Cell 9, 773–788 (2002).

2. Haering, C.H. et al. Structure and stability of cohesin’s Smc1-kleisin interaction. Mol Cell 15, 951–964 (2004).

3. Peters, J.M., Tedeschi, A. & Schmitz, J. The cohesin complex and its roles in chromosome biology. Genes Dev 22, 3089–3114 (2008).

4. Gruber, S., Haering, C.H. & Nasmyth, K. Chromosomal cohesin forms a ring. Cell 112, 765–777 (2003).

5. Yatskevich, S., Rhodes, J. & Nasmyth, K. Organization of Chromosomal DNA by SMC Complexes. Annu Rev Genet 53, 445–482 (2019).

6. Haering, C.H., Farcas, A.M., Arumugam, P., Metson, J. & Nasmyth, K. The cohesin ring concatenates sister DNA molecules. Nature 454, 297–301 (2008).

7. Srinivasan, M. et al. The Cohesin Ring Uses Its Hinge to Organize DNA Using Non-topological as well as Topological Mechanisms. Cell 173, 1508–1519 e1518 (2018).

8. Hauf, S., Waizenegger, I.C. & Peters, J.M. Cohesin cleavage by separase required for anaphase and cytokinesis in human cells. Science 293, 1320–1323 (2001).

9. Tachibana-Konwalski, K. et al. Rec8-containing cohesin maintains bivalents without turnover during the growing phase of mouse oocytes. Genes Dev 24, 2505–2516 (2010).

10. Uhlmann, F., Lottspeich, F. & Nasmyth, K. Sister-chromatid separation at anaphase onset is promoted by cleavage of the cohesin subunit Scc1. Nature 400, 37–42 (1999).

11. Uhlmann, F., Wernic, D., Poupart, M.A., Koonin, E.V. & Nasmyth, K. Cleavage of cohesin by the CD clan protease separin triggers anaphase in yeast. Cell 103, 375–386 (2000).

12. Fudenberg, G., Abdennur, N., Imakaev, M., Goloborodko, A. & Mirny, L.A. Emerging Evidence of Chromosome Folding by Loop Extrusion. Cold Spring Harb Symp Quant Biol 82, 45–55 (2017).

13. Fudenberg, G. et al. Formation of Chromosomal Domains by Loop Extrusion. Cell Rep 15, 2038–2049 (2016).

14. Sanborn, A.L. et al. Chromatin extrusion explains key features of loop and domain formation in wild-type and engineered genomes. Proc Natl Acad Sci U S A 112, E6456–6465 (2015).

15. Davidson, I.F. et al. DNA loop extrusion by human cohesin. Science 366, 1338–1345 (2019).

16. Kim, Y., Shi, Z., Zhang, H., Finkelstein, I.J. & Yu, H. Human cohesin compacts DNA by loop extrusion. Science 366, 1345–1349 (2019).

17. Golfier, S., Quail, T., Kimura, H. & Brugues, J. Cohesin and condensin extrude DNA loops in a cell cycle-dependent manner. Elife 9 (2020).

18. Rao, S.S. et al. A 3D map of the human genome at kilobase resolution reveals principles of chromatin looping. Cell 159, 1665–1680 (2014).

19. de Wit, E. et al. CTCF Binding Polarity Determines Chromatin Looping. Mol Cell 60, 676–684 (2015).

20. Guo, Y. et al. CRISPR Inversion of CTCF Sites Alters Genome Topology and Enhancer/Promoter Function. Cell 162, 900–910 (2015).

21. Vietri Rudan, M. et al. Comparative Hi-C reveals that CTCF underlies evolution of chromosomal domain architecture. Cell Rep 10, 1297–1309 (2015).

22. Nora, E.P. et al. Targeted Degradation of CTCF Decouples Local Insulation of Chromosome Domains from Genomic Compartmentalization. Cell 169, 930–944 e922 (2017).

23. Wutz, G. et al. Topologically associating domains and chromatin loops depend on cohesin and are regulated by CTCF, WAPL, and PDS5 proteins. EMBO J 36, 3573–3599 (2017).

24. Li, Y. et al. The structural basis for cohesin-CTCF-anchored loops. Nature 578, 472–476 (2020).

25. Rao, S.S.P. et al. Cohesin Loss Eliminates All Loop Domains. Cell 171, 305–320 e324 (2017).

26. Crane, E. et al. Condensin-driven remodelling of X chromosome topology during dosage compensation. Nature 523, 240–244 (2015).

27. Zuin, J. et al. Cohesin and CTCF differentially affect chromatin architecture and gene expression in human cells. Proc Natl Acad Sci U S A 111, 996–1001 (2014).

28. Bintu, B. et al. Super-resolution chromatin tracing reveals domains and cooperative interactions in single cells. Science 362 (2018).

29. Gibcus, J.H. et al. A pathway for mitotic chromosome formation. Science 359 (2018).

30. Naumova, N. et al. Organization of the mitotic chromosome. Science 342, 948–953 (2013).

31. Oomen, M.E., Hansen, A.S., Liu, Y., Darzacq, X. & Dekker, J. CTCF sites display cell cycle-dependent dynamics in factor binding and nucleosome positioning. Genome Res 29, 236–249 (2019).

32. Gassler, J. et al. A mechanism of cohesin-dependent loop extrusion organizes zygotic genome architecture. EMBO J 36, 3600–3618 (2017).

33. Abramo, K. et al. A chromosome folding intermediate at the condensin-to-cohesin transition during telophase. Nat Cell Biol 21, 1393–1402 (2019).

34. Schwarzer, W. et al. Two independent modes of chromatin organization revealed by cohesin removal. Nature 551, 51–56 (2017).

35. Belaghzal, H. et al. Liquid chromatin Hi-C characterizes compartment-dependent chromatin interaction dynamics. Nat Genet 53, 367–378 (2021).

36. Shi, Z., Gao, H., Bai, X.C. & Yu, H. Cryo-EM structure of the human cohesin-NIPBL-DNA complex. Science (2020).

37. Mitter, M. et al. Conformation of sister chromatids in the replicated human genome. Nature 586, 139–144 (2020).

38. Widom, J. Physicochemical studies of the folding of the 100 A nucleosome filament into the 300 A filament. Cation dependence. J Mol Biol 190, 411–424 (1986).

39. Sanders, J.T. et al. Loops, TADs, Compartments, and Territories are Elastic and Robust to Dramatic Nuclear Volume Swelling. bioRxiv (2021).

40. Dekker, J. Mapping in vivo chromatin interactions in yeast suggests an extended chromatin fiber with regional variation in compaction. J Biol Chem 283, 34532–34540 (2008).

41. Borrie, M.S., Campor, J.S., Joshi, H. & Gartenberg, M.R. Binding, sliding, and function of cohesin during transcriptional activation. Proc Natl Acad Sci U S A 114, E1062–E1071 (2017).

42. Busslinger, G.A. et al. Cohesin is positioned in mammalian genomes by transcription, CTCF and Wapl. Nature 544, 503–507 (2017).

43. Lengronne, A. et al. Cohesin relocation from sites of chromosomal loading to places of convergent transcription. Nature 430, 573–578 (2004).

44. Higashi, T.L. et al. A Structure-Based Mechanism for DNA Entry into the Cohesin Ring. Mol Cell 79, 917–933 e919 (2020).

45. Higashi, T.L., Pobegalov, G., Tang, M., Molodtsov, M.I. & Uhlmann, F. A Brownian ratchet model for DNA loop extrusion by the cohesin complex. Elife 10, 2021.2002.2014.431132 (2021).

46. Natsume, T., Kiyomitsu, T., Saga, Y. & Kanemaki, M.T. Rapid Protein Depletion in Human Cells by Auxin-Inducible Degron Tagging with Short Homology Donors. Cell Rep 15, 210–218 (2016).

47. Ran, F.A. et al. Genome engineering using the CRISPR-Cas9 system. Nat Protoc 8, 2281–2308 (2013).

48. Pauli, A. et al. Cell-type-specific TEV protease cleavage reveals cohesin functions in Drosophila neurons. Dev Cell 14, 239–251 (2008).

49. Chu, V.T. et al. Increasing the efficiency of homology-directed repair for CRISPR-Cas9-induced precise gene editing in mammalian cells. Nat Biotechnol 33, 543–548 (2015).

50. Belaghzal, H., Dekker, J. & Gibcus, J.H. Hi-C 2.0: An optimized Hi-C procedure for high-resolution genome-wide mapping of chromosome conformation. Methods 123, 56–65 (2017).

51. Imakaev, M. et al. Iterative correction of Hi-C data reveals hallmarks of chromosome organization. Nat Methods 9, 999–1003 (2012).

52. Valton, A.-L. et al. A cohesin traffic pattern genetically linked to gene regulation. bioRxiv (2021).

53. Heinz, S. et al. Simple combinations of lineage-determining transcription factors prime cis-regulatory elements required for macrophage and B cell identities. Mol Cell 38, 576–589 (2010).

54. Ramirez, F. et al. deepTools2: a next generation web server for deep-sequencing data analysis. Nucleic Acids Res 44, W160–165 (2016).

55. Campagne, A. et al. BAP1 complex promotes transcription by opposing PRC1-mediated H2A ubiquitylation. Nat Commun 10, 348 (2019).

56. McHaourab, Z.F., Perreault, A.A. & Venters, B.J. ChIP-seq and ChIP-exo profiling of Pol II, H2A.Z, and H3K4me3 in human K562 cells. Sci Data 5, 180030 (2018).

