## Extended Data Figures for "Biochemically distinct cohesin complexes mediate positioned loops between CTCF sites and dynamic loops within chromatin domains"

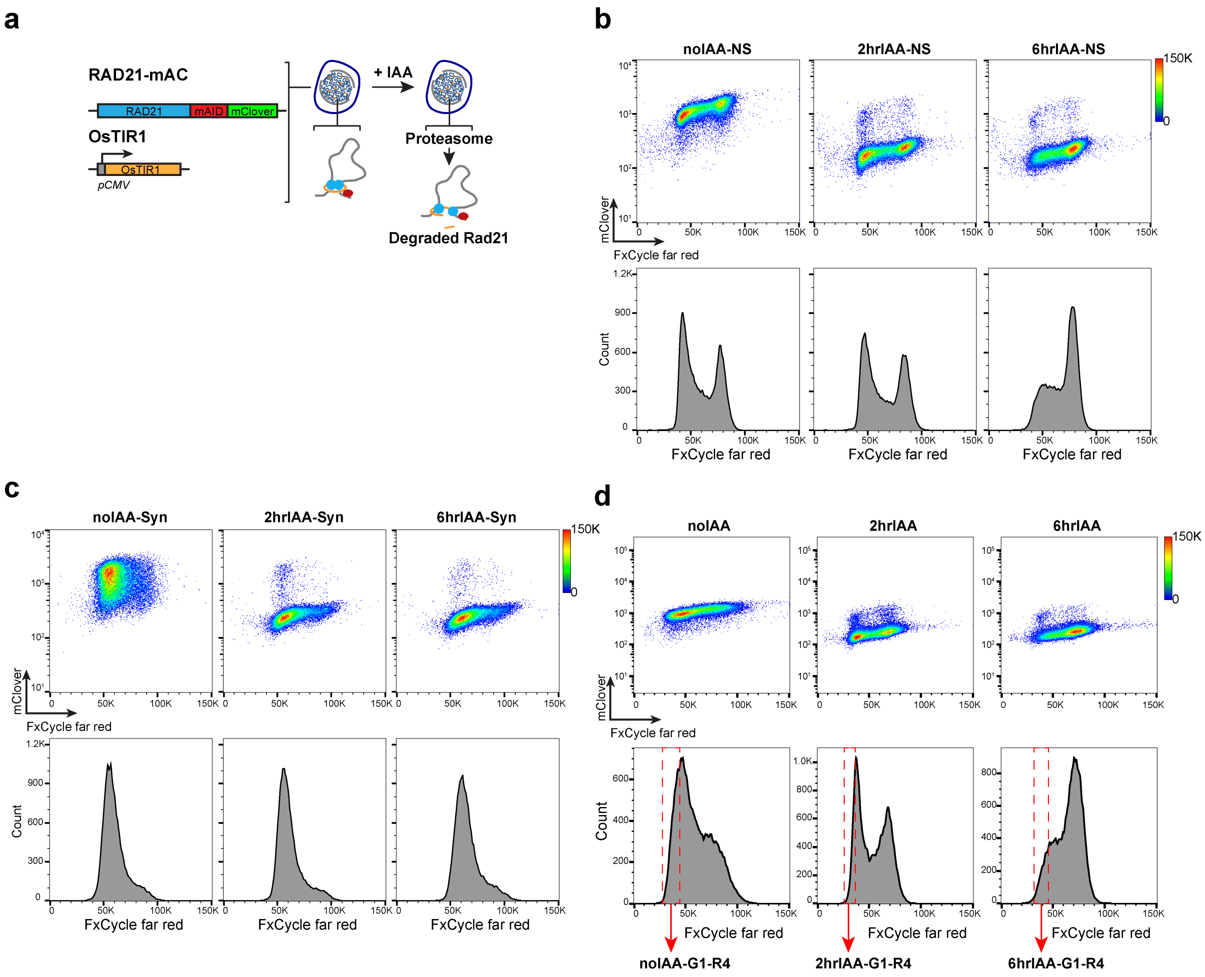


**Extended Data Fig. 1. Cell cycle profiles of the HCT-RAD21-mAC cells in different conditions for Hi-C experiments**

(**a)** Schematic of HCT-RAD21-mAC cells. The cell line was a gift from Dr. Kanemaki lab. AID sequence and mClover were inserted at the C-terminus of RAD21 and OsTIR with pCMV promoter were inserted into AAVS1 save harbor as previous described [46]. After treatments with 500uM IAA, RAD21-mAC was degraded. **(b)** FACS analysis of non-synchronous HCT116-mAC cells without and with 500uM IAA treatments for 2 and 6 hour. DNA was stained using fxCycle-far red and RAD21 was tagged with mClover. Upper panels are 2D scatter plots indicating DNA contents and RAD21 levels before and after IAA treatment. Lower panels, histograms indicating cell cycle stage distributions for cultures with and without IAA treatment. The same staining and sorting method was used for **(c)** and **(d)**. **(c)** FACS analysis of G1 synchronized HCT116-mAC cells without and with 500uM IAA treatment for 2 and 6. For G1 synchronization, HCT116-mAC cells were first treated with 2mM thymidine for 12 hours, then released in fresh medium for 12 hours, followed by treatment for 12 hours with 400uM mimosine[25]. Cells were arrested at the G1/S boundary. The medium of G1/S arrested cells as then replaced with medium + 400uM mimosine (untreated) or medium + 500uM IAA + 400uM mimosine for 2 or 6 hours. The cells were then fixed for FACS (lower panels) and Hi-C analysis. Upper panels: plots of DNA contents vs RAD21 levels with and without IAA treatment. Lower panels: cell cycle stage distributions with and without after IAA treatment. **(d)** FACS analysis of non-synchronous HCT116-mAC cells without and with 500uM IAA treatment for 2 and 6 hour treatments. G1 cells were sorted from these non-synchronous cells for Hi-C analysis. Upper panels are 2D scatter plots DNA content and RAD21 levels with and without IAA treatment. Lower panels, histogram graphics indicate cell cycle profiles with and without IAA treatment. Red dashed boxes indicate the G1 population that were sorted and collected for Hi-C analysis of G1 cells.

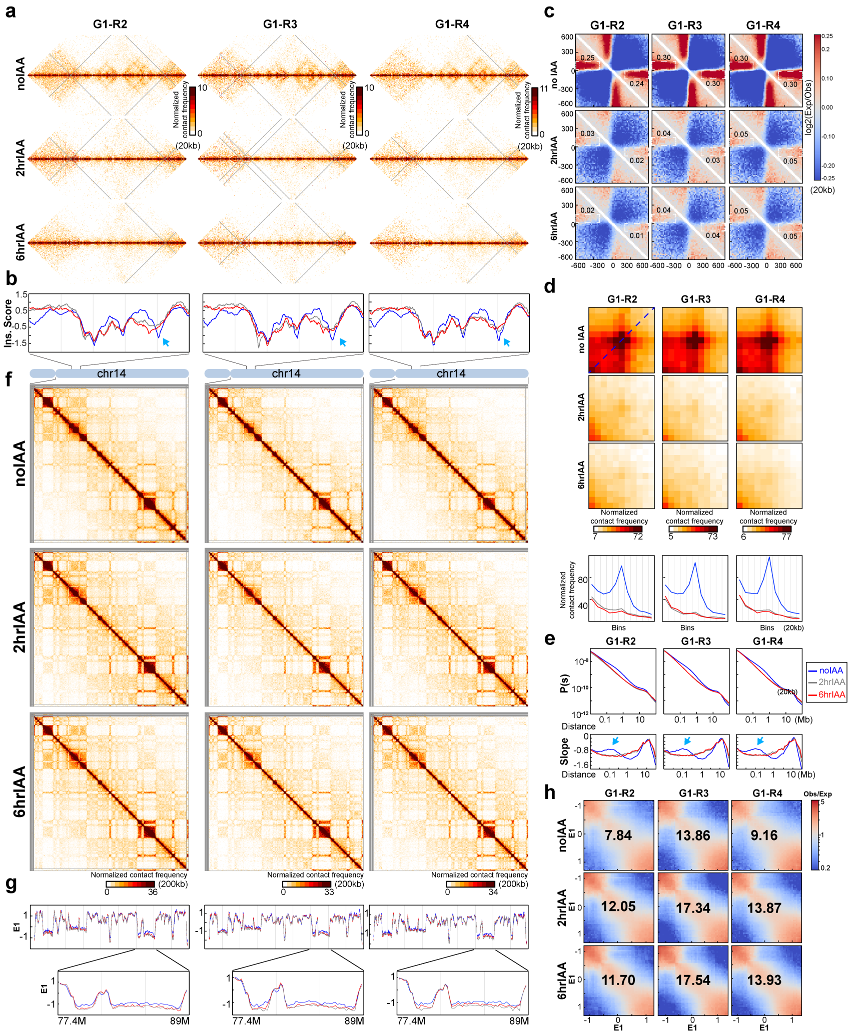

**Extended Data Fig. 2. Three replicates of Hi-C analysis of G1 cells without and with IAA treatment**

**(a)** Hi-C interaction maps for three independent Hi-C experiments using G1-sorted cells without and with IAA treatment for 2 and 6 hours respectively. Data for the 29-34 Mb regions of chromosome 14 is shown at 20kb resolution. **(b)** Insulation profiles for the 29-34 Mb regions of chromosome 14 at 20kb resolution. The blue, grey and red lines represent no IAA treatments, 2hour IAA and 6hour IAA treatments, respectively. Blue arrow shows weakened insulation at boundaries **(c)**. Aggregate Hi-C data binned at 20kb resolution at TAD boundaries that were identified in each replicate without IAA treatments. The numbers of TAD boundaries for each replicate are 4178, 4419 and 4362, respectively. The numbers at the sides of the cross indicate the strength of boundary-anchored stripes using the mean values of interaction frequency within the white dashed boxes. These values are related to boundary strength. **(d)**. Aggregated Hi-C data at a set of loops identified in HCT116-RAD21-mAC cells with intact RAD21 (n=3169) identified by [25]. Plots at the bottom show average Hi-C signals along the dotted blue lines representing signals from the bottom-left corner to the top-right corner of the loop aggregated heatmaps shown in upper panels. **(e)**. *P(s)* plots (upper panels) and the derivative from *P(s)* plots (lower panels) for Hi-C data as indicated. The arrows on the derivative plots indicate cohesin loops. **(f)**. Hi-C interaction maps for three independent Hi-C experiments using G1-sorted cells without and with IAA treatment for 2 and 6 hours respectively. Data for the 18-107.3 Mb regions of chromosome 14 is shown at 200kb resolution. **(g)**. Eigenvector value E1 across the 18-107.3 Mb regions of chromosome 14 at 200kb resolution. Bottom panels, E1 for the 77.4-89Mb region is shown at 200kb resolution and the color assignments for the lines are the same as for the other panels. **(h)**. Saddle plots of Hi-C data binned at 200kb for three independent Hi-C experiments using G1-sorted cells without and with IAA treatment for 2 and 6 hours respectively. Saddle plots for each cell condition were calculated using the E1 calculated from the Hi-C data obtained with G1 cells grown without IAA treatments. The numbers indicate compartment strength.

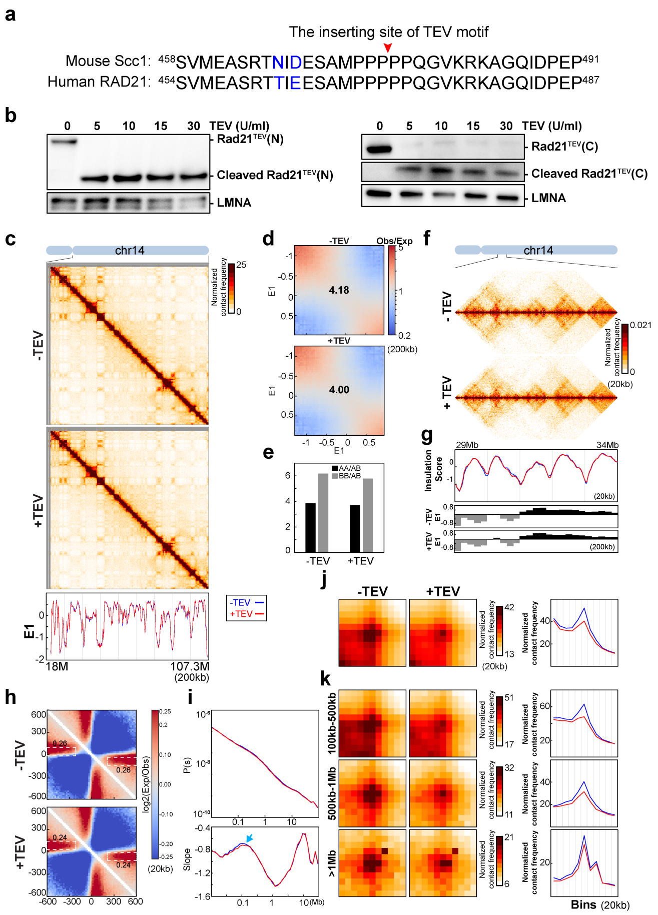


**Extended Data Fig. 3. A replicate Hi-C analysis of nuclei with RAD21 cleaved in NB buffer**

**(a)** The TEV motif inserting site is conserved between human and mouse. The amino acid sequences flanking TEV motif inserting sites in mouse Scc1 and human RAD21 are shown. **(b)** TEV proteases added at different concentrations cleaved RAD21 in purified nuclei from HAP1-RAD21$^{TEV}$. Left panels, RAD21$^{TEV}$ and cleaved fragments were detected using an antibody recognizing N-terminus of RAD21 $^{TEV}$ (Abcam ab154769) and LMNA was used as a loading control. Right panels, RAD21$^{TEV}$ and cleaved fragments were detected using an antibody recognizing C-terminus of RAD21$^{TEV}$ (Abcam ab992) and LMNA was used as a loading control. Concentrations of TEV proteases were indicated. **(c)**. Hi-C interaction maps for HAP1-RAD21$^{TEV}$ nuclei without and with TEV treatment, respectively. Data for the 18-107.3 Mb region of chromosome 14 is shown at 200kb resolution. Bottom panel, eigenvector E1 across the 18-107.3 Mb regions of chromosome 14 at 200kb resolution. **(d)**. Saddle plots of Hi-C data binned at 200kb for HAP1-RAD21$^{TEV}$ nuclei without and with TEV treatment, respectively. The numbers indicate compartment strength. **(e).** Interaction strength of compartments. The bars represent the strength of compartment interactions for each sample as indicated. Dark bars, strength of AA interaction as compared to AB interaction (A-A/A-B). Grey bars, strength of BB interaction as compared to AB interaction (B-B/B-A). **(f)**. Hi-C interaction maps for HAP1-RAD21$^{TEV}$ nuclei without and with TEV treatment, respectively. Data for the 29-34 Mb region of chromosome 14 is shown at 20kb resolution. **(g)**. Insulation profiles for the 29-34 Mb regions of chromosome 14 at 20kb resolution. The blue and red lines represent without and with TEV protease treatment, respectively. The lower panels indicate compartment Eigenvector value E1 across the same region at 200kb resolution. **(h).** Aggregate Hi-C data binned at 20kb resolution at TAD boundaries identified in the sample in NB buffer without TEV treatment. The numbers at the sides of the cross indicate the strength of boundary-anchored stripes using the mean values of interaction frequency within the white dashed boxes, as in Fig. 1e. **(i)** *P(s)* plots (upper panels), and the derivatives of *P(s)* plots (lower panels) for Hi-C data from nuclei with or without TEV treatment as indicated. The blue arrows indicate the signature of cohesin loops in each condition. **(j)** Aggregated Hi-C data binned at 20kb resolution at loops as in Fig. 3j. Right panel: average Hi-C signals along the blue dashed line shown in the left Hi-C panel. **(k)** Aggregated Hi-C data binned at 20kb resolution at chromatin loops of three different loop sizes, 100-500kb, 500kb-1Mb, and >1Mb. Right panels: average Hi-C signals along the blue dashed line shown in the left Hi-C map in panel Fig. 3j.

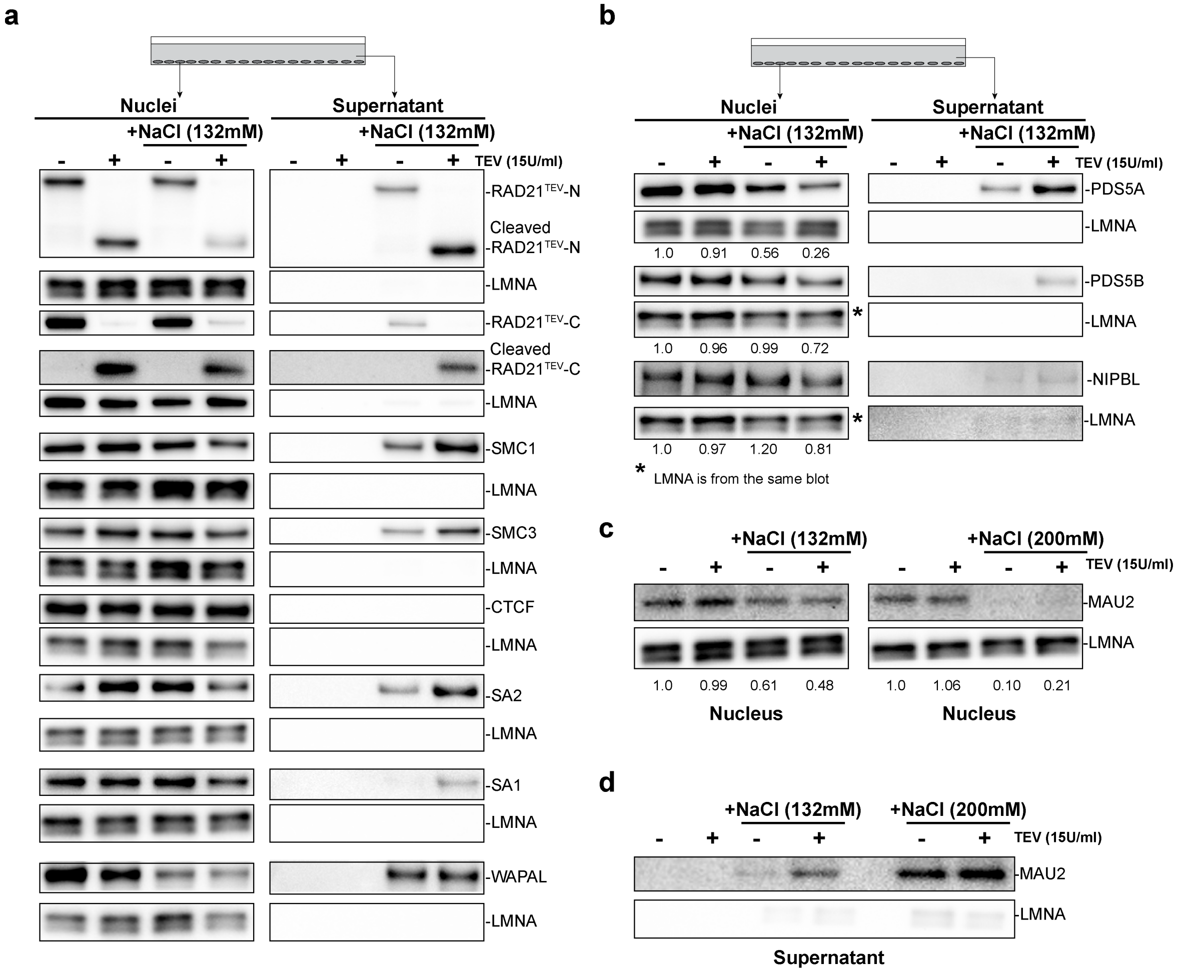


**Extended Data Fig. 4. Cleaving RAD21 in NBS1 dissociates and releases cohesin components**

Western blot analysis of nuclear retention of cohesin in RAD21$^{TEV}$ nuclei with and without TEV protease treatments in low salt (NB) or salt (NBS1) buffers. Nuclei were purified from HAP1-RAD21$^{TEV}$ cells and plated on poly-Lys plates in 4ml NB buffer or NB buffer with 132mM NaCl. As indicated, 15U/ml of TEV protease was added to cleave RAD21 and all the plates were kept at 4°C for overnight. Left panels indicated as nuclei show cohesin components in nuclei detected using antibodies as described in Figure 4a and 4b. Right panels indicated as supernatant analyze cohesin components released to the buffer: 3ml buffer was collected from the plates after overnight incubation and the buffer was spun at 800g for 10 minutes to remove any unattached nuclei. 2ml of the supernatant was then concentrated using the Amicon column (3kd cut-off) at 12,000g to obtain the volume of around 100ul (20X). The concentrated supernatant was adjusted to 100ul and transferred to a new tube with 25ul 5X loading sample buffer, then boiled for western analysis. Cohesin components from supernatants were separated and detected using the same antibodies as used for the Western blots shown on the left Figure S4abc. For all western blot analyses, LMNA was used as the loading controls. (**a**) A biological replicate of analysis of cohesin subunits retained in nuclei ore released to the supernatant with and without TEV treatment in NB or NBS1 buffers. The antibodies used here are the same as Fig. 4a. (**b**) Western blot analysis of PDS5A, PDS5B and NIPBL retained in nuclei or released to the supernatant with and without TEV treatment in NB or NBS1 buffers. (**c**) and (**d**) Western analysis of MAU2 (SCC4) retained in nuclei (c) and released to the supernatant (d) with and without TEV treatment in NB, NBS1 and NBS2 buffers.

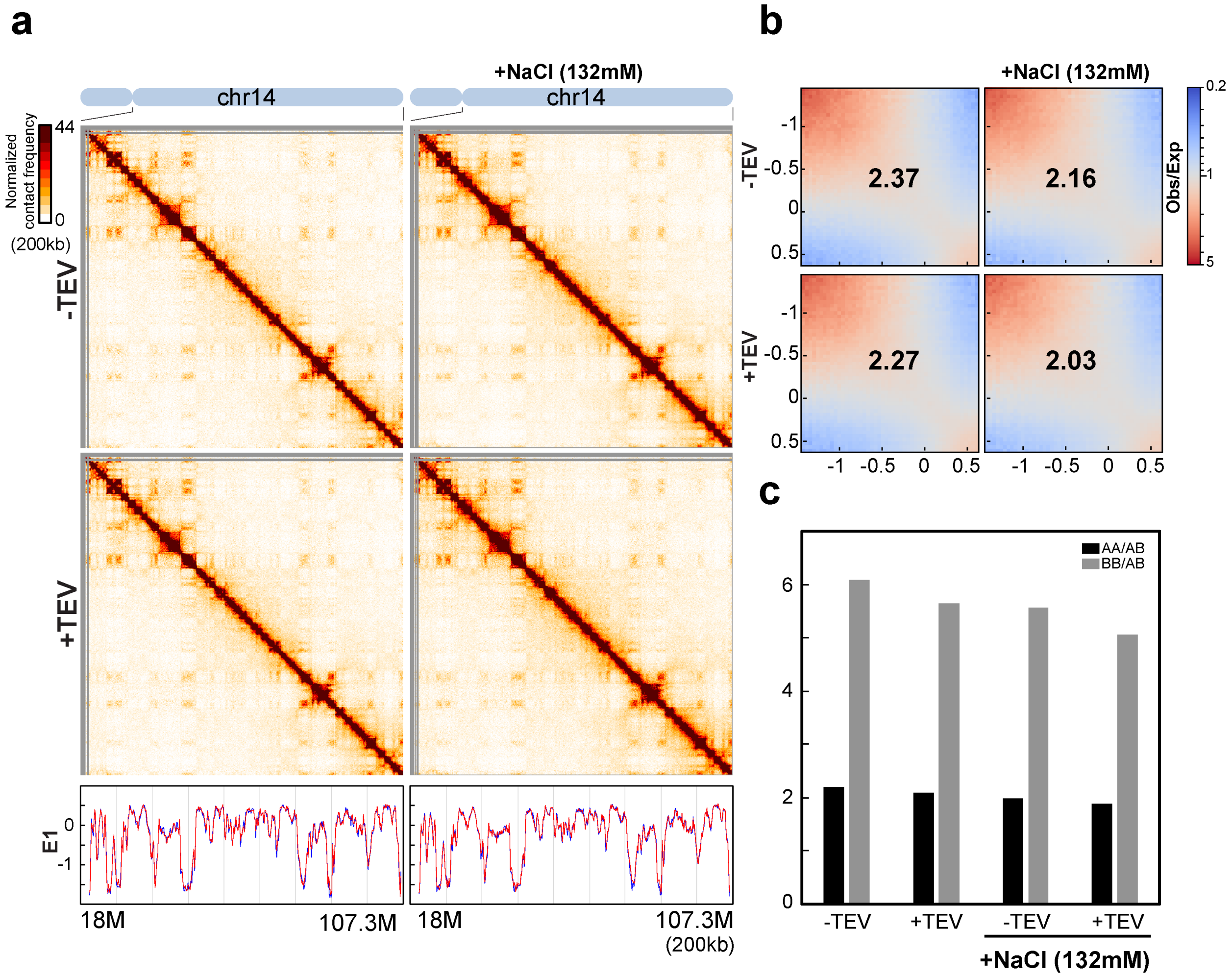


**Extended Data Fig. 5. Cleaving RAD21 in NBS1 has little effects on compartmentalization and CTCF binding** **(a)** Hi-C interaction maps for HAP1-RAD21$^{TEV}$ nuclei without and with TEV treatment, respectively. Data for the 18-107.3 Mb region of chromosome 14 is shown at 200kb resolution. Bottom, eigenvector E1 profiles across the 18-107.3 Mb region of chromosome 14 at 200kb resolution for each sample. **(b)** Saddle plots of Hi-C data binned at 200kb for HAP1-RAD21$^{TEV}$ nuclei without and with TEV treatment, respectively. The numbers indicate compartment strength. **(c)** Interaction strength of compartments. The bars represent the strength of compartment interactions for each sample as indicated. Dark bars, strength of AA interaction as compared to AB interaction (A-A/A-B). Grey bars, strength of BB interaction as compared to AB interaction (B-B/B-A).

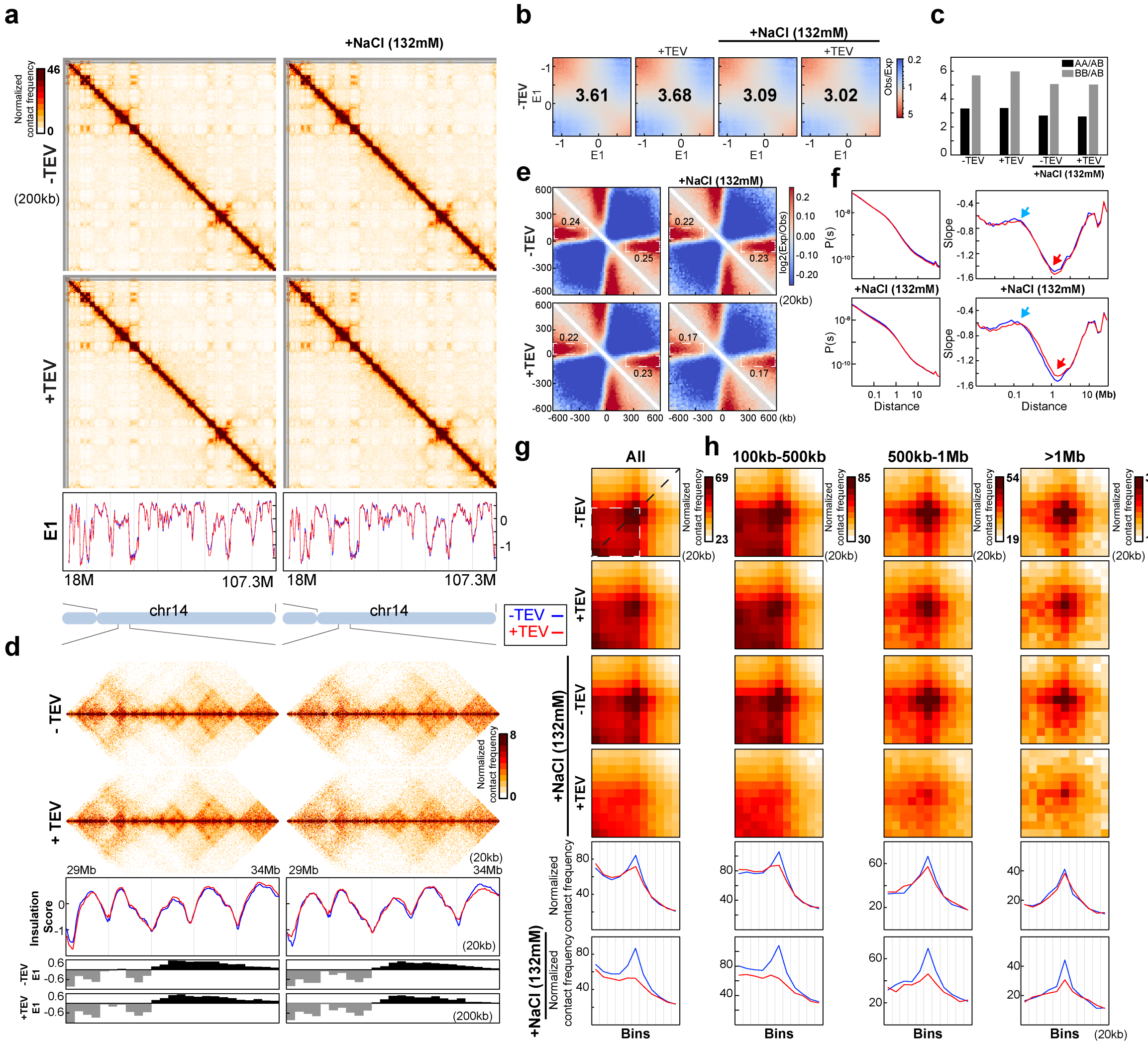

**Extended Data Fig. 6. A replicate Hi-C analysis of the nuclei with cleaved RAD21 in NBS1**

**(a)** Hi-C interaction maps for HAP1-RAD21$^{TEV}$ nuclei without and with TEV protease treatment in NB or NBS1, respectively. Data for the 18-107.3 Mb region of chromosome 14 is shown at 200kb resolution. Right panels, 132mM NaCl was added to NB buffer during TEV protease treatments at 4°C overnight. Bottom, eigenvector E1 across the 18-107.3 Mb region of chromosome 14 at 200kb resolution. **(b)** Saddle plots of Hi-C data binned at 200kb for HAP1-RAD21$^{TEV}$ nuclei without and with TEV treatment in NB or NBS1, respectively. The numbers indicate compartment strength. **(c)** Interaction strength of compartments. The bars represent the strength of compartment interactions for each sample as indicated. Dark bars, strength of AA interaction as compared to AB interaction (A-A/A-B). Grey bars, strength of BB interaction as compared to AB interaction (B-B/B-A). **(d)** Hi-C interaction maps for salt effects on HAP1-RAD21$^{TEV}$ nuclei without and with TEV protease treatment in NB or NBS1, respectively. Data for the 29-34 Mb regioncof chromosome 14 is shown at 20kb resolution. Middle panels, insulation profiles for the 29-34 Mb regions of chromosome 14 at 20kb resolution. The lower panels indicate compartment Eigenvector value E1 across the same region at 200kb resolution. **(e)** Aggregate Hi-C data binned at 20kb resolution at TAD boundaries identified in the sample in NB buffer without TEV treatment in NB or NBS1. The numbers at the sides of the cross indicate the strength of boundary-anchored stripes using the mean values of interaction frequency within the white dashed boxes, as in Fig. 1e. **(f)** *P(s)* plots (left panels), and the derivatives of *P(s)* plots (right panels) for Hi-C data from nuclei with or without TEV treatment as indicated. The blue arrows indicate the signature of cohesin loops in each condition. The red arrows indicate the changes of contact frequency at 2Mb. **(g)** Aggregated Hi-C data binned at 20kb resolution at loops as in Fig. 3i. Lower panels: average Hi-C signals along the blue dashed line shown in the upper left Hi-C panel. **(k)** Aggregated Hi-C data binned at 20kb resolution at chromatin loops of three different loop sizes, 100-500kb, 500kb-1Mb, and >1Mb. Lower panels:  average Hi-C signals along the blue dashed line shown in the left Hi-C map in panel Fig. 3j.

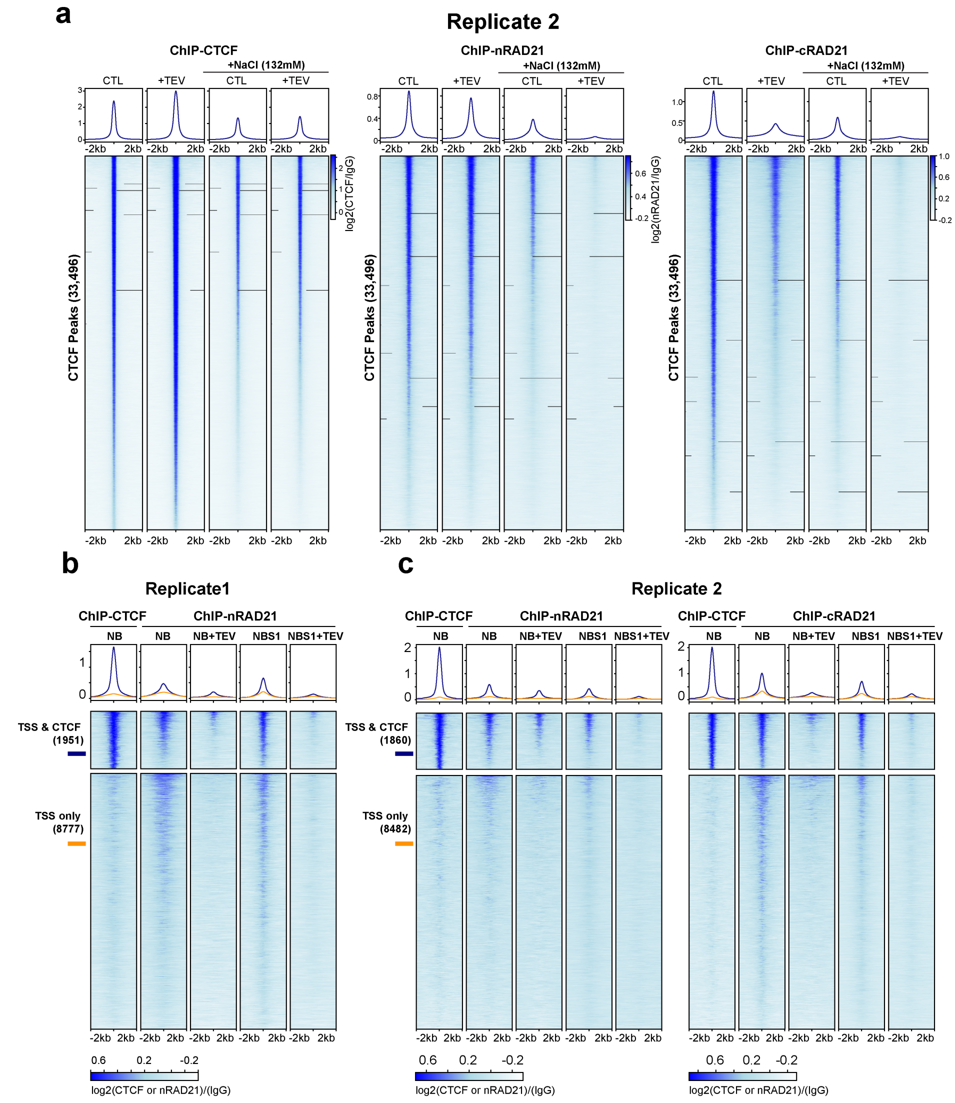


**Extended Data Fig. 7. A replicate of ChIP analysis of the nuclei with cleaved RAD21 in NBS1**

**(a)** Profiles of ChIP signals of CTCF and RAD21 at the CTCF binding sites without and with TEV treatment in NB or NBS1 buffer. Data shown are for biological replicate 2, a independent replicate of the experiment shown in Fig. 4h (Extended Table 3). 33,496 CTCF binding sites were identified from the CTCF ChIP data of nuclei without TEV treatment in NB. Upper panel, the average CTCF and RAD21 ChIP-seq signals for each condition for the set of 33,496 CTCF binding sites. Two different RAD21 antibodies were used, nRAD21 is ab154 from Abcam recognizing the N-terminal domain of RAD21, as used in Figure 4h. The cRAD21 is ab992 from Abcam recognizing C-terminal domain of RAD21. Lower panel, stack-up heatmap of CTCF and RAD21 ChIP-seq signals of each condition at each of the 33,496 CTCF binding sites. **(b)** Profiles of ChIP signals of CTCF and RAD21 at active transcription start sites (TSS) without and with TEV in NB or NBS1 buffer. Of 13,412 active TSS of HAP1 cells, 1951 overlapped with 30,148 CTCF binding sites while 8777 did not overlap with CTCF binding sites (2kb away from CTCF binding sites). Both average ChIP signals (upper panels) and stack-up heatmap of ChIP signals of CTCF and RAD21 for these two groups of TSS sites are shown. **(c)** A biological replicate of the experiment shown in panel b. Of 13412 active TSS sites, 1860 were overlapped with 33,496 CTCF binding sites while 8482 did not overlap with CTCF binding sites. Dark blue and orange lines indicate TSSs that overlapped or did not overlap with CTCF binding sites, respectively. Both average ChIP signals (upper panels) and stack-up heatmap of ChIP signals of CTCF and RAD21 on these two groups of TSS sites are shown. Right panel includes a RAD21 ChIP data using the RAD21 antibody that recognizes C-terminal of RAD21. Dark blue and orange lines indicate TSS that overlapped or did not overlap with CTCF binding sites, respectively.

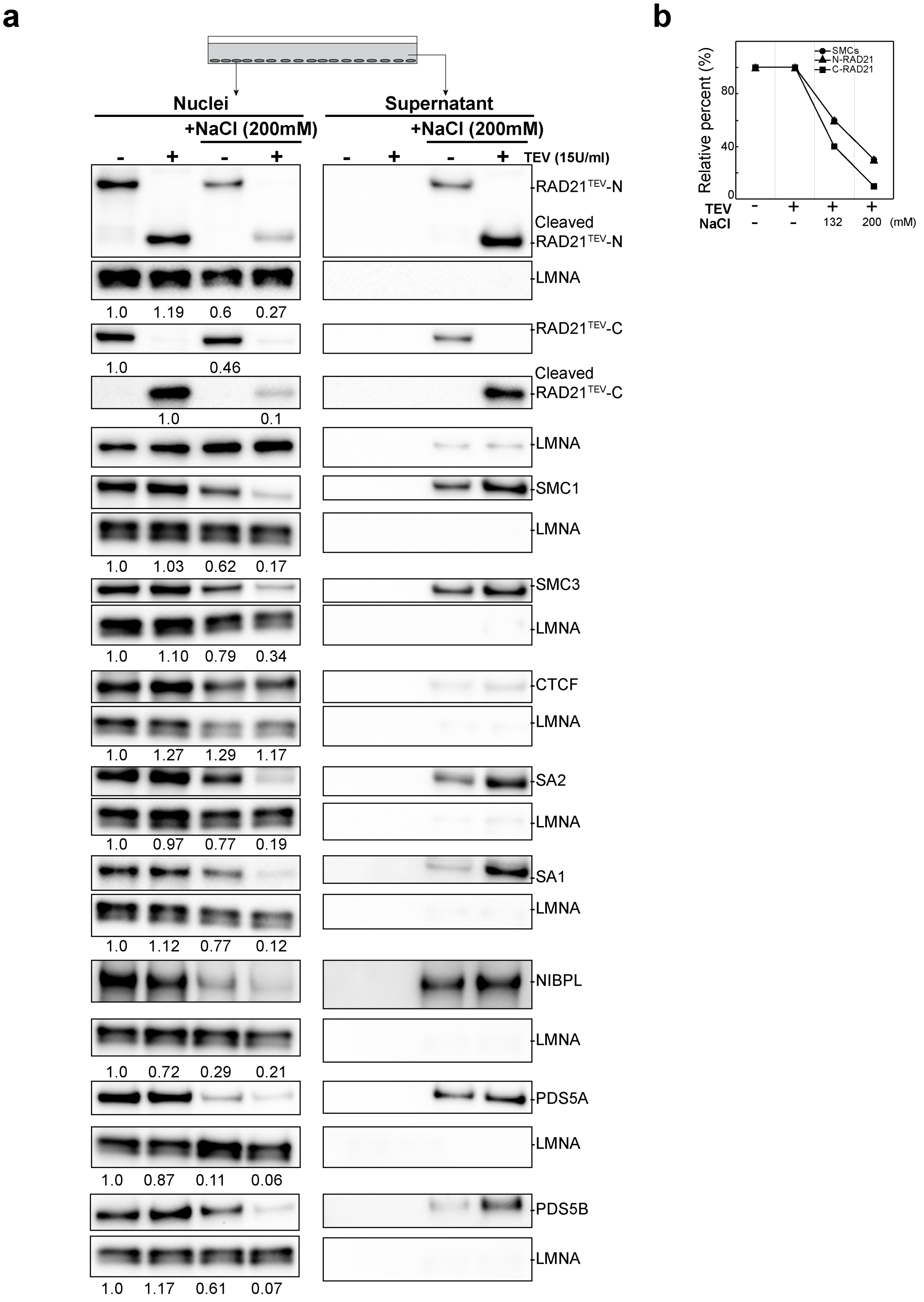


**Extended Data Fig. 8. Cleaving RAD21 in NBS2 dissociates and releases cohesin components**

**(a)** Western blot analysis of salt effects on nuclear retention of cohesin complex subunits in HAP-1RAD21$^{TEV}$ nuclei with and without TEV protease treatment. Western blot analysis was performed as indicated in Figure S4a except that 200mM NaCl was used, instead of 132mM. **(b)** Quantification of levels of SMC1/3, N-terminal cleaved RAD21 and C-terminal cleaved RAD21. The levels of indicated proteins or fragments were normalized to LMNA as loading controls first, then the ratio was normalized to the same protein or fragments in NB without TEV treatment. For SMC proteins, the mean relative percentage of SMC1 and SMC3 lecels, from Figure 4a and Extended Data Fig. 8, is presented.

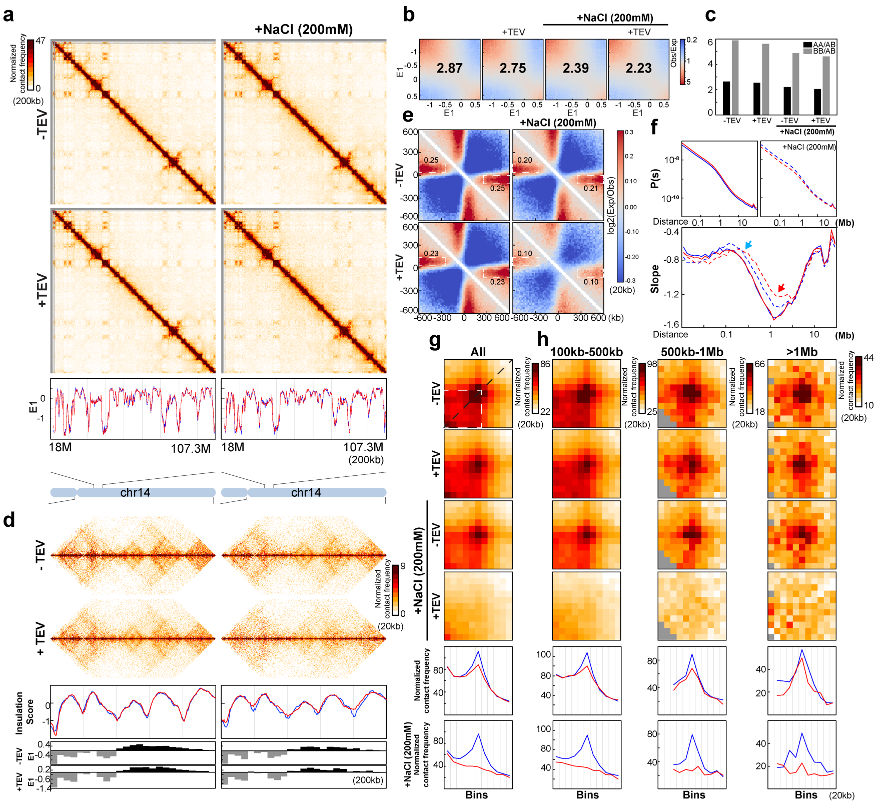


Extended Data Fig. S9

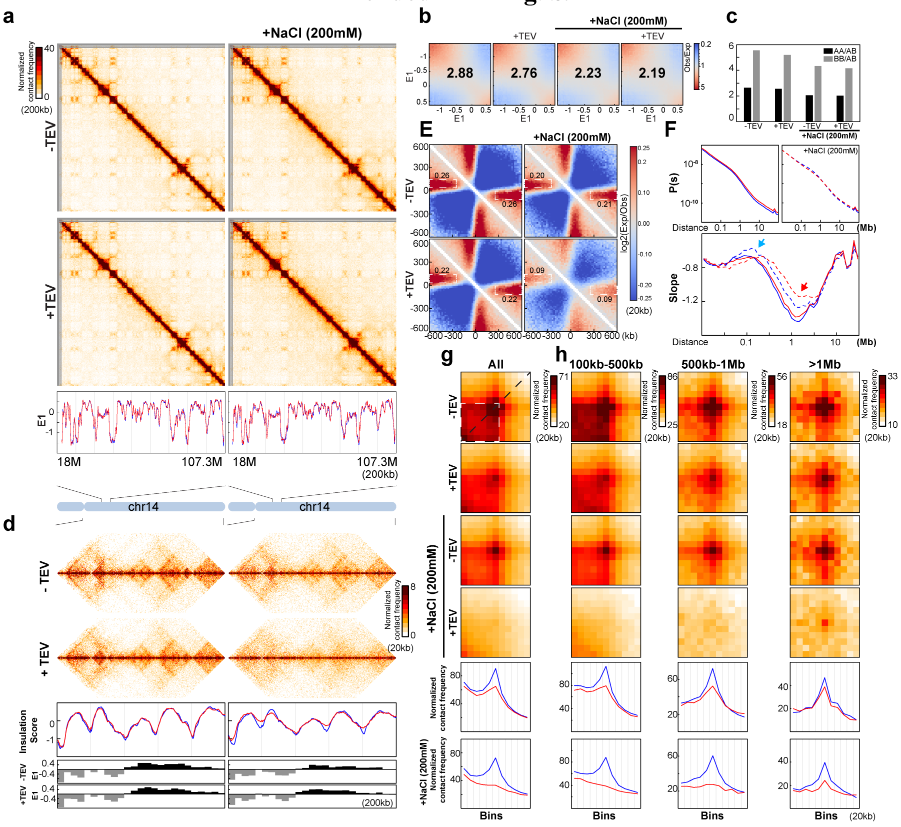


Extended Data Fig. 10

**Extended Data Fig. S9-10. Two biological replicates of Hi-C analysis of nuclei with RAD21 cleaved in NBS2 buffer**

**(a)** Hi-C interaction maps for HAP1-RAD21$^{TEV}$ nuclei without and with TEV protease treatments in NB or NBS2 (NB with 200mM NaCl), respectively. Data for the 18-107.3 Mb region of chromosome 14 is shown at 200kb resolution. Bottom, eigenvector E1 across the 18-107.3 Mb region of chromosome 14 at 200kb resolution. **(b)** Saddle plots of Hi-C data binned at 200kb for HAP1-RAD21$^{TEV}$ nuclei without and with TEV treatments in NB or NBS2 buffer, respectively. The numbers indicate compartment strength. **(c)** Interaction strength of compartments. The bars represent the strength of compartment interactions for each sample as indicated. Dark bars, strength of AA interaction as compared to AB interaction (A-A/A-B). Grey bars, strength of BB interaction as compared to AB interaction (B-B/B-A). **(d)** Hi-C interaction maps for HAP1-RAD21$^{TEV}$ nuclei without and with TEV treatment in NB or NBS2 buffer, respectively. Data for the 29-34 Mb region of chromosome 14 is shown at 20kb resolution. Middle panels indicate insulation profiles for the 29-34 Mb regions of chromosome 14 at 20kb resolution. The blue and red lines represent without and with TEV protease treatment, respectively, as in panel a. The lower panels indicate compartment Eigenvector value E1 across the same region at 200kb resolution. **(e)** Aggregate Hi-C data binned at 20kb resolution at TAD boundaries identified in the sample in NB buffer without TEV treatment. The numbers at the sides of the cross indicate the strength of boundary-anchored stripes using the mean values of interaction frequency within the white dashed boxes, as in Fig. 1e. **(f)** *P(s)* plots (left panels), and the derivatives of *P(s)* plots (right panels) for Hi-C data from nuclei with or without TEV treatment as indicated. The blue arrows indicate the signature of cohesin loops in each condition. The red arrows indicate the changes of contact frequency at 2Mb. (**g**) Aggregated Hi-C data binned at 20kb resolution at loops as in Fig. 3j. Lower panels: average Hi-C signals along the blue dashed line shown in the upper left Hi-C panel. **(h)** Aggregated Hi-C data binned at 20kb resolution at chromatin loops of three different loop sizes, 100-500kb, 500kb-1Mb, and >1Mb. Lower panels: average Hi-C signals along the blue dashed line shown in the left Hi-C map in Fig. panel 3j.

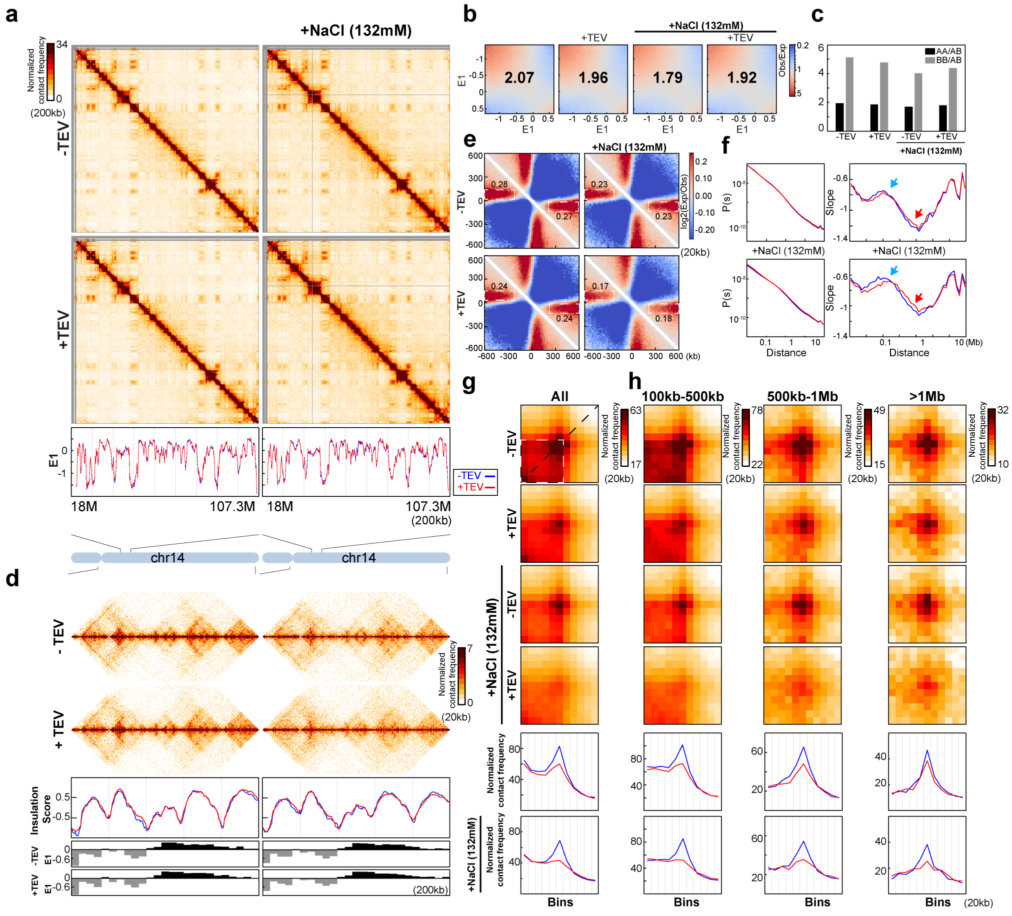


**Extended Data Fig. 11**

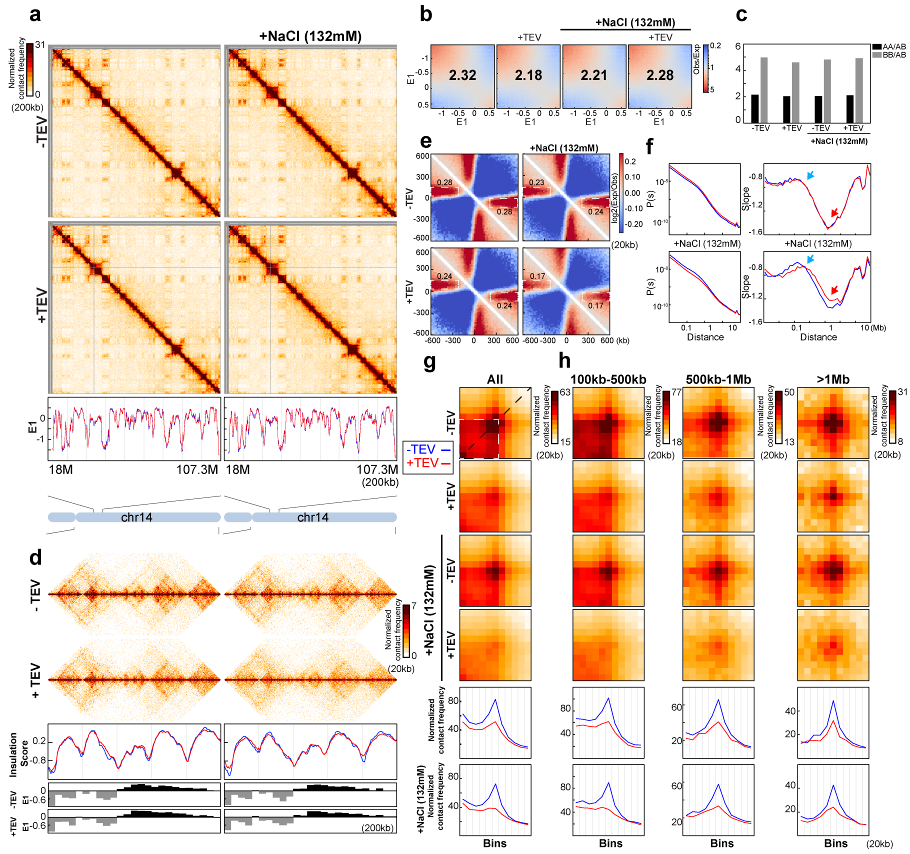

**Extended Data Fig. 12**

**Extended Data Fig. 11-12. Two biological replicates of Hi-C analysis of G1-sorted nuclei with RAD21 cleaved in NBS1.**

To perform Hi-C analysis on sorted G1 nuclei, around 20 million nuclei were incubated overnight in 12ml indicated buffers in 15ml tubes without and with TEV (15U/ml) at 4C. After treatment, nuclei were fixed using 1% FA and then stained with PI for sorting G1 nuclei similarly as previously described for HCT-116-RAD21-mAC cells. Around 1 million of G1 nuclei were obtained after sorting for subsequent Hi-C experiments. **(a)** Hi-C interaction maps for G1-sorted HAP1-RAD21$^{TEV}$ nuclei without and with TEV protease treatment in NB or NBS1 buffer, respectively. Data for the 18-107.3 Mb region of chromosome 14 is shown at 200kb resolution. Bottom, eigenvector E1 cross the 18-107.3 Mb region of chromosome 14 at 200kb resolution. **(b)** Saddle plots of Hi-C data binned at 200kb for G1-sorted HAP1-RAD21$^{TEV}$ nuclei without and with TEV treatment in NB or NBS1 buffer, respectively. The numbers indicate compartment strength. **(c)** Interaction strength of compartments. The bars represent the strength of compartment interactions for each sample as indicated. Dark bars, strength of AA interaction as compared to AB interaction (A-A/A-B). Grey bars, strength of BB interaction as compared to AB interaction (B-B/B-A). **(d)** Hi-C interaction maps for G1-sorted HAP1-RAD21$^{TEV}$ nuclei without and with TEV treatment in NB or NBS1 buffer, respectively. Data for the 29-34 Mb region of chromosome 14 is shown at 20kb resolution. Middle panels indicate insulation profiles for the 29-34 Mb regions of chromosome 14 at 20kb resolution. The blue and red lines represent without and with TEV protease treatment, respectively, as in panel a. The lower panels indicate compartment Eigenvector value E1 across the same region at 200kb resolution. **(e)** Aggregate Hi-C data binned at 20kb resolution at TAD boundaries identified in the sample in NB buffer without TEV treatment. The numbers at the sides of the cross indicate the strength of boundary-anchored stripes using the mean values of interaction frequency within the white dashed boxes, as in Fig. 1e. **(f)** *P(s)* plots (left panels), and the derivatives of *P(s)* plots (right panels) for Hi-C data from nuclei with or without TEV treatment as indicated. The blue arrows indicate the signature of cohesin loops in each condition. The red arrows indicate the changes of contact frequency at 2Mb. (**g**) Aggregated Hi-C data binned at 20kb resolution at loops as in Fig. 3j. Lower panel: average Hi-C signals along the blue dashed line shown in the left Hi-C panel. **(h)** Aggregated Hi-C data binned at 20kb resolution at chromatin loops of three different loop sizes, 100-500kb, 500kb-1Mb, and >1Mb. Lower panels: average Hi-C signals along the blue dashed line shown in the left Hi-C map in panel Fig. 3j.

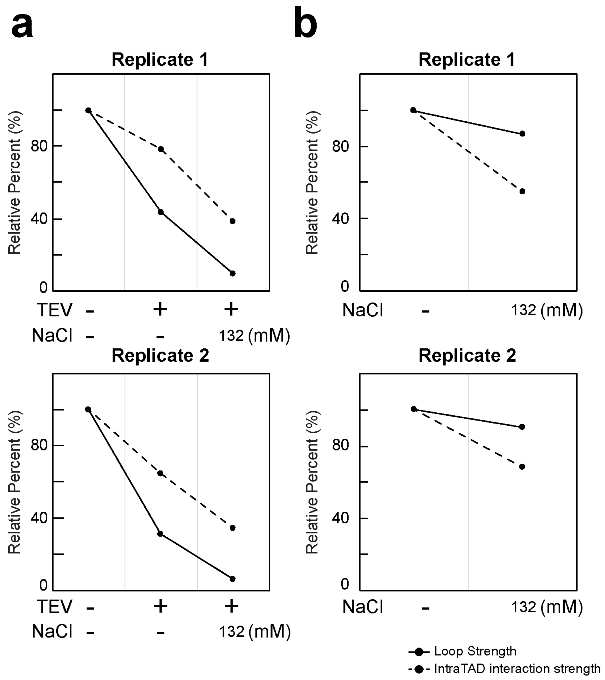


**Extended Data Fig. 13. Quantification of loop and intra-TAD analysis for two biological replicates of the G1 nuclei with RAD21 cleaved in NBS1**

**(a)** Quantification of loop strength and intra-TAD interaction strength obtained with G1-sorted HAP1-RAD21$^{TEV}$ nuclei without and with TEV treatment in NB, NBS1 buffer, respectively. Loop strength and intra-TAD interaction strength were calculated as in Fig. 1g. Loop strength and intra-TAD interaction strength from nuclei in NB without TEV treatment was used to normalize respectively. Intra-TAD interaction strength was further normalized by the baseline level of 36%, observed in the complete absence of cohesin, and that reflects the general distance-dependent decay of interaction frequency as shown in Fig. 1g. Two biological replicates are shown. **(b)** Quantification of loop strength and intra-TAD interaction strength without TEV treatment in NB or NBS1 buffer, respectively. Loop strength and intra-TAD interaction strength were normalized as in Fig. 4i.

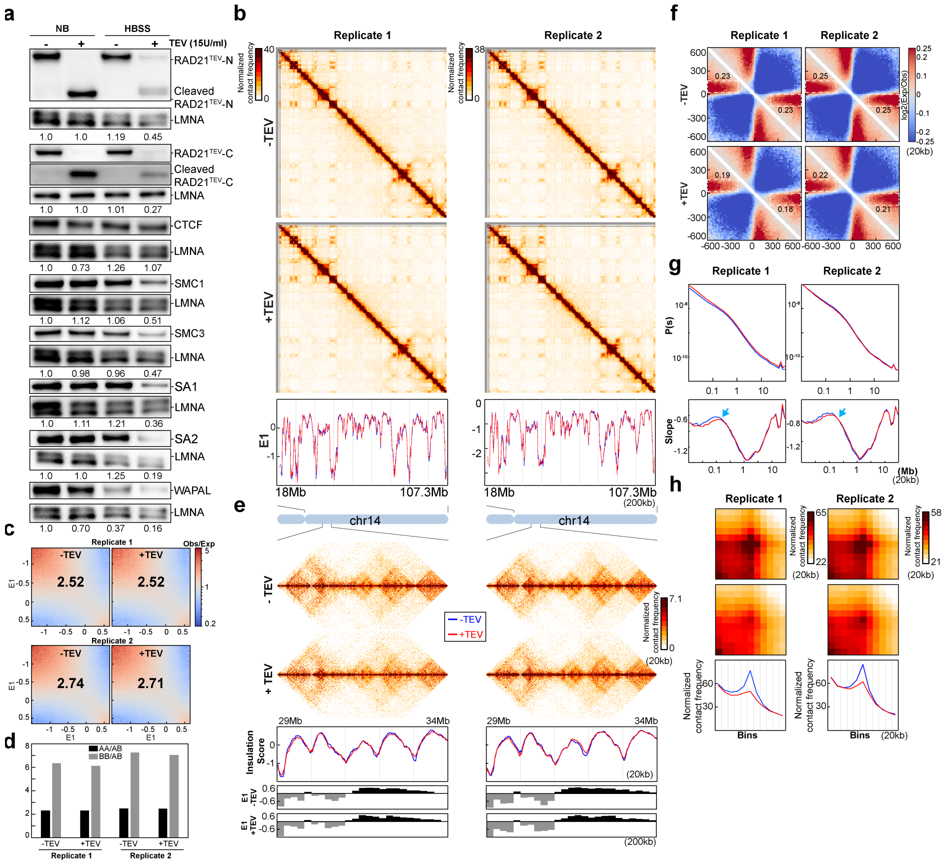


**Extended Data Fig. 14. Two replicates of Hi-C analysis of the nuclei with RAD21 cleaved in HBSS buffer**

**(a)** Western blot analysis of RAD21 and cohesin components with and without TEV protease treatments in NB and HBSS buffer, respectively. The purified nuclei from HAP1-RAD21$^{TEV}$ were plated in NB and HBSS buffer, respectively, and incubated without and with TEV proteases at 4°C overnight. The nuclei were then lysed and cohesin components were separated on gels and detected using antibodies as shown (See methods for all the antibody information for this analysis). LMNA was used as loading controls and levels of each cohesin subunit in each condition was normalized first to LMNA, then to that of nuclei in NB without TEV protease treatment to obtain relative levels. **(b)** Hi-C interaction maps for HAP1-RAD21$^{TEV}$ nuclei without and with TEV protease treatment in HBSS buffer, respectively. Data for the 18-107.3 Mb region of chromosome 14 is shown at 200kb resolution. Bottom, eigenvector E1 cross the 18-107.3 Mb region of chromosome 14 at 200kb resolution. **(c)** Saddle plots of Hi-C data binned at 200kb for HAP1-RAD21$^{TEV}$ nuclei without and with TEV treatments in HBSS buffer, respectively. The numbers indicate compartment strength. **(d)** Interaction strength of compartments. The bars represent the strength of compartment interactions for each sample as indicated. Dark bars, strength of AA interaction as compared to AB interaction (A-A/A-B). Grey bars, strength of BB interaction as compared to AB interaction (B-B/B-A). **(e)** Hi-C interaction maps for HAP1-RAD21$^{TEV}$ nuclei without and with TEV treatment in HBSS buffer, respectively. Data for the 29-34 Mb region of chromosome 14 is shown at 20kb resolution. Middle panels indicate insulation profiles for the 29-34 Mb regions of chromosome 14 at 20kb resolution. The blue and red lines represent without and with TEV protease treatments, respectively, as in panel b. The lower panels indicate compartment Eigenvector value E1 across the same region at 200kb resolution. **(f)** Aggregate Hi-C data binned at 20kb resolution at TAD boundaries identified in the sample without TEV treatment in each replicate. The numbers at the sides of the cross indicate the strength of boundary-anchored stripes using the mean values of interaction frequency within the white dashed boxes, as in Fig. 1e. **(g)** *P(s)* plots (upper panels), and the derivatives of *P(s)* plots (lower panels) for Hi-C data from nuclei with or without TEV treatment as indicated. The blue arrows indicate the signature of cohesin loops in each condition. (**h**) Aggregated Hi-C data binned at 20kb resolution at loops as in Fig. 3j. Lower panel: average Hi-C signals along the blue dashed line shown in the left Hi-C panel.

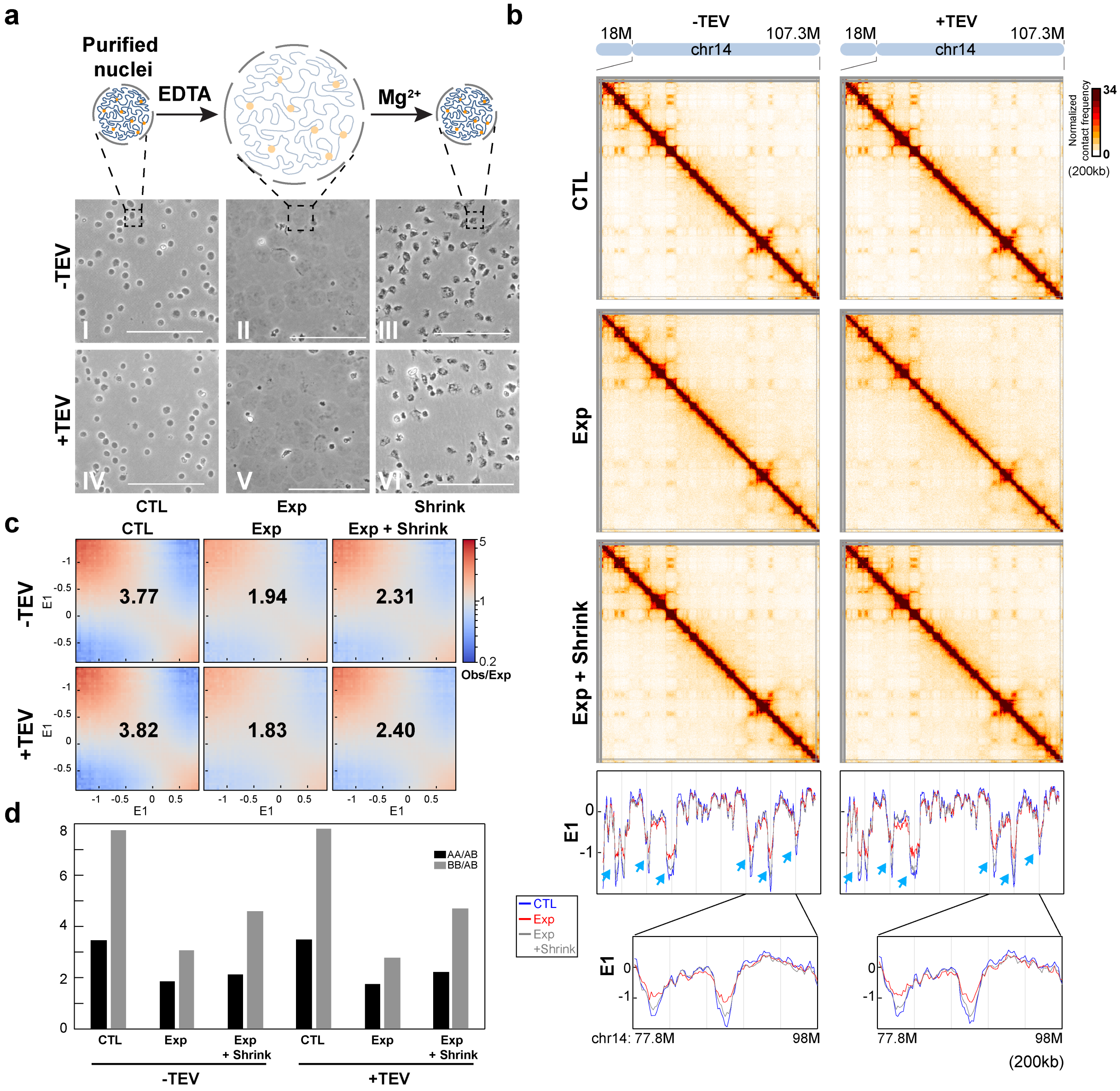


**Extended Data Fig. 15. A biological replicate for compartmentalization analysis after nuclear expansion and contraction**

**(a)** Changes of nuclear morphology during the expansion and contraction assay. Upper panel: a schematic to show nuclear morphology changes before expansion, after expansion, and after expansion followed by compaction. Middle and bottom, images of HAP1-RAD21$^{TEV}$ nuclei before expansion, after expansion, and after expansion followed by compaction without and with TEV protease treatment, respectively (Scalebar =100μm). **(b)** Hi-C interaction maps for HAP1-RAD21$^{TEV}$ nuclei before expansion, after expansion, and after expansion followed by compaction without and with TEV protease treatment, respectively. Data for the 18-107.3 Mb region of chromosome 14 is shown at 200kb resolution. Lower panels: Eigenvector E1 across the 18-107.3 Mb region of chromosome 14 at 200kb resolution. The blue, red and grey lines represent before expansion, after expansion, and after expansion followed by compaction treatments, respectively. Blue arrows indicate E1 changes. Bottom panels, E1 across the 77.8-98Mb region of chromosome 14 at 200kb resolution. **(c)** Saddle plots of Hi-C data binned at 200kb for RAD21$^{TEV}$ nuclei before expansion, after expansion, and after expansion followed by compaction without and with TEV protease treatment, respectively. The numbers indicate compartment strength. **(d)** Interaction strength of compartments. The bars represent the strength of compartment interactions for each sample as indicated. Dark bars, strength of AA interaction as compared to AB interaction (A-A/A-B). Grey bars, strength of BB interaction as compared to AB interaction (B-B/B-A).

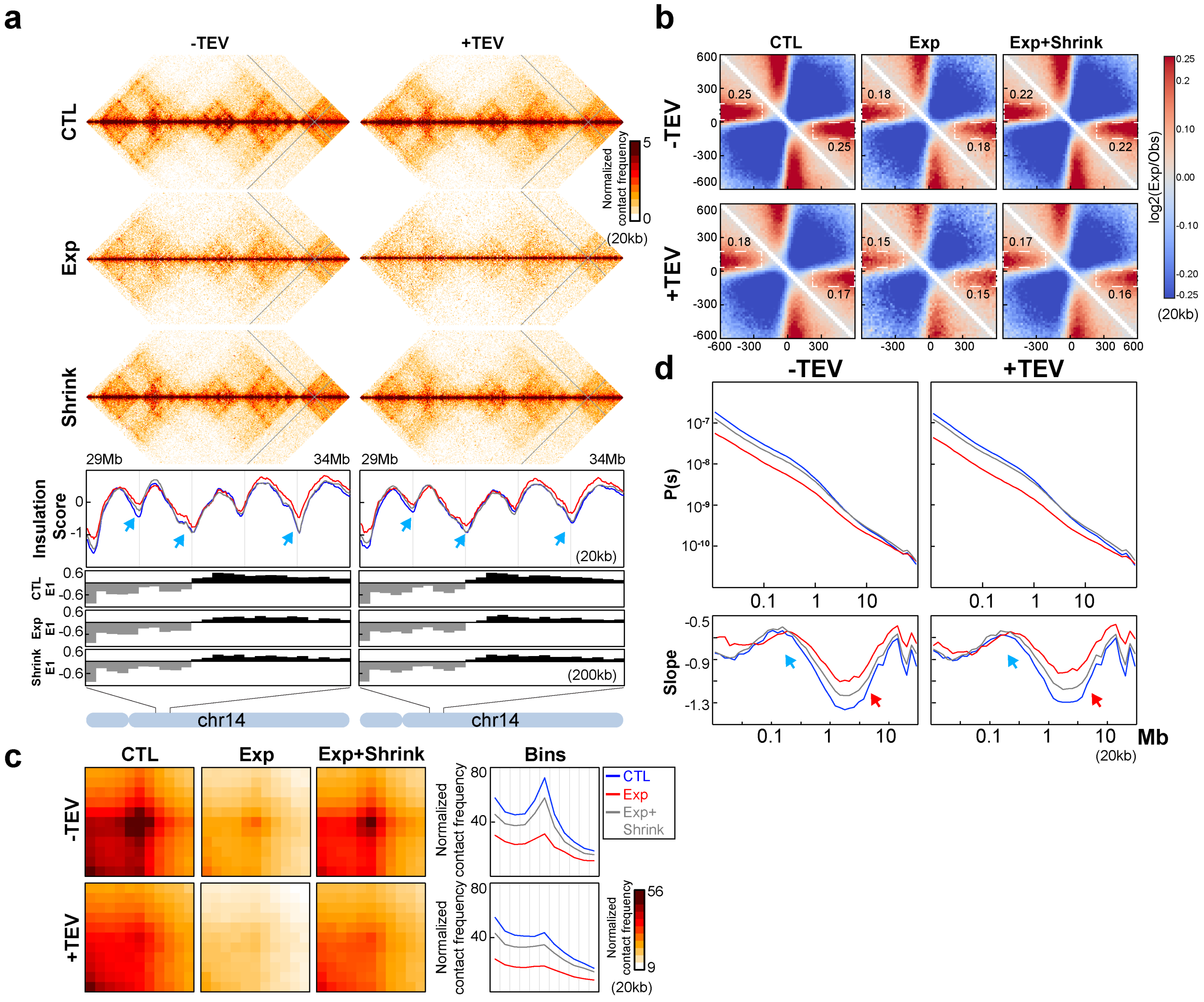


**Extended Data Fig. 16. A biological replicate for Hi-C analysis after nucleus expansion and contraction**

**(a)** Hi-C interaction maps for HAP1-RAD21$^{TEV}$ nuclei before expansion, after expansion, and after expansion followed by compaction without and with TEV protease treatment, respectively. Data for the 29-34 Mb region of chromosome 14 is shown at 20kb resolution. Insulation profiles for the 29-34 Mb regions of chromosome 14 at 20kb resolution. The blue, red and grey lines represent before expansion, after expansion, and after expansion followed by compaction treatments, respectively. The blue arrows indicate weakened insulation boundary as nuclei was expanded. The lower panels indicate compartment Eigenvector value E1 across the same region at 200kb resolution. **(b)** Aggregate Hi-C data binned at 20kb resolution at TAD boundaries identified in control sample without TEV treatment. The numbers at the sides of the cross indicate the strength of boundary-anchored stripes using the mean values of interaction frequency within the white dashed boxes, as in Fig. 1e. (**c**) Aggregated Hi-C data binned at 20kb resolution at loops as used in Fig. 3j. Right panels: average Hi-C signals along the blue dashed line shown in the left upper Hi-C panel. **(d)** *P(s)* plots (upper panels), and the derivatives of *P(s)* plots (lower panels) for Hi-C data from nuclei before expansion, after expansion, and after expansion followed by compaction without and with TEV protease treatment, as indicated. The blue arrows indicate the signature of cohesin loops in each condition. The red arrows indicate the changes of contact frequency at 2Mb.

| Antibodies | Source | Cat. Number | Usage |
|---|---|---|---|
| Rabbit RAD21, N-terminal | Abcam | ab154769 | WB, ChIP |
| Rabbit RAD21, C-terminal | Abcam | ab992 | WB, ChIP |
| Rabbit Lamin A | Abcam | ab26300 | WB |
| Rabbit SMC3 | Abcam | ab9263 | WB |
| Rabbit SMC1 | Bethyl lab | A300-055A | WB |
| Rabbit CTCF | Millipore | 07-729 | WB, ChIP |
| Rabbit SA2 | Bethyl lab | A302-580A | WB |
| Rabbit SA1 | Bethyl lab | A302-579A | WB |
| Rabbit WAPAL | Cell Signaling Tech | 77428 | WB |
| Rabbit NIPBL | Bethyl lab | A301-778A | WB |
| Rabbit SCC-112 (PDS5A) | Bethyl lab | A300-089A | WB |
| Rabbit PDS5B | Bethyl lab | A300-538A | WB |
| Rabbit Mau2 (SCC4) antibody | Novus Biologics | NBP1-56779 | WB |

**Extended Data Table 1. All the antibodies used in this study.**

| Hi-C Experiments | Hi-C librares | Cis | Trans | Total uniqe mapped | cis% | Related Figures |
|---|---|---|---|---|---|---|
| NS cells, noIAA | noIAA-ns-R1 | 76508241 | 9564906 | 86073147 | 88.89% | Fig. 1 and 2 |
| NS cells, 2hrIAA | 2hrIAA-ns-R1 | 65861487 | 12125631 | 77987118 | 84.45% | Fig. 1 and 2 |
| NS cells, 6hrIAA | 6hrIAA-ns-R1 | 67255389 | 12933948 | 80189337 | 83.87% | Fig. 1 and 2 |
| Syn cells, noIAA | noIAA-syn-R1 | 77480896 | 10650654 | 88131550 | 87.92% | Fig. 1 and 2 |
| Syn cells, 2hrIAA | 2hrIAA-syn-R1 | 62043810 | 13181327 | 75225137 | 82.48% | Fig. 1 and 2 |
| Syn cells, 6hrIAA | 6hrIAA-syn-R1 | 69720083 | 13485549 | 83205632 | 83.79% | Fig. 1 and 2 |
| G1 cells, noIAA | noIAA-G1 | 101654170 | 20189530 | 121843700 | 83.43% | Fig. 1 and 2 |
| G1 cells, 2hrIAA | 2hrIAA-G1 | 81368607 | 20050134 | 101418741 | 80.23% | Fig. 1 and 2 |
| G1 cells, 6hrIAA | 6hrIAA-G1 | 86768371 | 25376248 | 112144619 | 77.37% | Fig. 1 and 2 |
| G1 cells, noIAA-R2 | noIAA-G1-R2 | 36364905 | 8268812 | 44633717 | 81.47% | Extended Fig. 2 |
| G1 cells, noIAA-R3 | noIAA-G1-R3 | 31407868 | 4167321 | 35575189 | 88.29% | Extended Fig. 2 |
| G1 cells, noIAA-R4 | noIAA-G1-R4 | 33881397 | 7753397 | 41634794 | 81.38% | Extended Fig. 2 |
| G1 cells, 2hrIAA-R2 | 2hrIAA-G1-R2 | 25072229 | 6787217 | 31859446 | 78.70% | Extended Fig. 2 |
| G1 cells, 2hrIAA-R3 | 2hrIAA-G1-R3 | 27904172 | 5423185 | 33327357 | 83.73% | Extended Fig. 2 |
| G1 cells, 2hrIAA-R4 | 2hrIAA-G1-R4 | 28392206 | 7839732 | 36231938 | 78.36% | Extended Fig. 2 |
| G1 cells, 6hrIAA-R2 | 6hrIAA-G1-R2 | 29682592 | 9299262 | 38981854 | 76.14% | Extended Fig. 2 |
| G1 cells, 6hrIAA-R3 | 6hrIAA-G1-R3 | 26972315 | 6212728 | 33185043 | 81.28% | Extended Fig. 2 |
| G1 cells, 6hrIAA-R4 | 6hrIAA-G1-R4 | 30113464 | 9864258 | 39977722 | 75.33% | Extended Fig. 2 |
| RAD21TEV, NB, noTEV, R2 | YL-CTL-NB-R2 | 86968698 | 95097319 | 182066017 | 47.77% | Fig. 3 |
| RAD21TEV, NB, TEV, R2 | YL-TEVON-NB-R2 | 86015138 | 89217000 | 175232138 | 49.09% | Fig. 3 |
| RAD21TEV, NB, noTEV, R1 | YL-CTL-NB-R1 | 108091965 | 25344581 | 133436546 | 81.01% | Extended Fig. 3 |
| RAD21TEV, NB, TEV, R1 | YL-TEVON-NB-R1 | 107470903 | 29522242 | 136993145 | 78.45% | Extended Fig. 3 |
| RAD21TEV, NB, noTEV, R6 | YL-CTL-NB-R6 | 41588558 | 38303876 | 79892434 | 52.06% | Fig.4 and Extended Fig. 5 |
| RAD21TEV, NB, TEV, R6 | YL-TEVON-NB-R6 | 48536267 | 47556326 | 96092593 | 50.51% | Fig.4 and Extended Fig. 5 |
| RAD21TEV, NB+132mM NaCl, noTEV, R6 | YL-CTL-NB-NaCl-R6 | 58982235 | 47168860 | 106151095 | 55.56% | Fig.4 and Extended Fig. 5 |
| RAD21TEV, NB+132mM NaCl, TEV, R6 | YL-TEVON-NaCl-R6 | 44949444 | 45593164 | 90542608 | 49.64% | Fig.4 and Extended Fig. 5 |
| RAD21TEV, NB, noTEV, R5 | YL-CTL-NB-R5 | 49281288 | 40246215 | 89527503 | 55.05% | Extended Fig. 6 |
| RAD21TEV, NB, TEV, R5 | YL-TEVON-NB-R5 | 46649455 | 36737096 | 83386551 | 55.94% | Extended Fig. 6 |
| RAD21TEV, NB+132mM NaCl, noTEV, R3 | YL-CTL-NB-NaCl-R3 | 37182546 | 33374197 | 70556743 | 52.70% | Extended Fig. 6 |
| RAD21TEV, NB+132mM NaCl, TEV, R3 | YL-TEVON-NaCl-R3 | 36088396 | 34596836 | 70685232 | 51.06% | Extended Fig. 6 |
| RAD21TEV, NB, noTEV, R8 | YL-NB-R8-T1 | 35738629 | 31814454 | 67553083 | 52.90% | Extended Fig. 9 |
| RAD21TEV, NB, TEV, R8 | YL-NB-TEV-R8-T1 | 44059977 | 36773746 | 80833723 | 54.51% | Extended Fig. 9 |
| RAD21TEV, NB+200mM NaCl, noTEV, R8 | YL-NB-salt-R8-T1 | 39858683 | 29002422 | 68861105 | 57.88% | Extended Fig. 9 |
| RAD21TEV, NB+200mM NaCl, TEV, R8 | YL-NB-salt-TEV-R8-T1 | 31690389 | 31432331 | 63122720 | 50.20% | Extended Fig. 9 |
| RAD21TEV, NB, noTEV, R10 | YL-NB-R10-T1 | 37670044 | 37019599 | 74689643 | 50.44% | Extended Fig. 10 |
| RAD21TEV, NB, TEV, R10 | YL-NB-TEV-R10-T1 | 46919674 | 46967207 | 93886881 | 49.97% | Extended Fig. 10 |
| RAD21TEV, NB+200mM NaCl, noTEV, R10 | YL-NBS2-R10-T1 | 43384288 | 40123815 | 83508103 | 51.95% | Extended Fig. 10 |
| RAD21TEV, NB+200mM NaCl, TEV, R10 | YL-NBS2-TEV-R10-T1 | 43707443 | 51017928 | 94725371 | 46.14% | Extended Fig. 10 |
| RAD21TEV, NB, noTEV, G1 sorted, R1 | YL-G1-NB-R1 | 49379378 | 42474628 | 91854006 | 53.76% | Extended Fig. 11 |
| RAD21TEV, NB, TEV, G1 sorted, R1 | YL-G1-NB-TEV-R1 | 52477105 | 48007000 | 100484105 | 52.22% | Extended Fig. 11 |
| RAD21TEV, NB+132mM NaCl, noTEV, G1 sorted, R1 | YL-G1-NB-salt-R1 | 48247579 | 65963200 | 114210779 | 42.24% | Extended Fig. 11 |
| RAD21TEV, NB+132mM NaCl, TEV, G1 sorted, R1 | YL-G1-NB-satl-TEV-R1 | 49147583 | 50167199 | 99314782 | 49.49% | Extended Fig. 11 |
| RAD21TEV, NB, noTEV, G1 sorted, R2 | YL-G1-NB-R2 | 49399975 | 47643815 | 97043790 | 50.90% | Extended Fig. 12 |
| RAD21TEV, NB, TEV, G1 sorted, R2 | YL-G1-NB-TEV-R2 | 47727105 | 40764159 | 88491264 | 53.93% | Extended Fig. 12 |
| RAD21TEV, NB+132mM NaCl, noTEV, G1 sorted, R2 | YL-G1-NB-salt-R2 | 50002144 | 37099573 | 87101717 | 57.41% | Extended Fig. 12 |
| RAD21TEV, NB+132mM NaCl, TEV, G1 sorted, R2 | YL-G1-NB-satl-TEV-R2 | 56835637 | 32267095 | 89102732 | 63.79% | Extended Fig. 12 |
| RAD21TEV, HBSS, noTEV, R4 | YL-NuCTL-DpnII-R4 | 44755703 | 39149121 | 83904824 | 53.34% | Extended Fig. 14 |
| RAD21TEV, HBSS, TEV, R4 | YL-NuTEVON-DpnII-R4 | 51845205 | 41033791 | 92878996 | 55.82% | Extended Fig. 14 |
| RAD21TEV, HBSS, noTEV, R5 | YL-NuCTL-DpnII-R5 | 56676056 | 34919294 | 91595350 | 61.88% | Extended Fig. 14 |
| RAD21TEV, HBSS, TEV, R5 | YL-NuTEVON-DpnII-R5 | 52183723 | 31809735 | 83993458 | 62.13% | Extended Fig. 14 |
| RAD21TEV, CTL, noTEV, R1 | YL-NuCTL4hr-R1 | 112229907 | 29223995 | 141453902 | 79.34% | Fig. 5 and 6 |
| RAD21TEV, Exp, noTEV, R1 | YL-NuExp4hr-R1 | 88914853 | 54256569 | 143171422 | 62.10% | Fig. 5 and 6 |
| RAD21TEV, Exp-Shrink, noTEV, R1 | YL-NuExp4hr-shrink-R1 | 98298632 | 34362901 | 132661533 | 74.10% | Fig. 5 and 6 |
| RAD21TEV, CTL, TEV, R1 | YL-NuCTL4hr-Tev-R1 | 102518642 | 30862442 | 133381084 | 76.86% | Fig. 5 and 6 |
| RAD21TEV, Exp, TEV, R1 | YL-NuExp4hr-Tev-R1 | 79125714 | 61468123 | 140593837 | 56.28% | Fig. 5 and 6 |
| RAD21TEV, Exp-Shrink, TEV, R1 | YL-NuExp4hr-shrink-Tev-R1 | 94890903 | 34570081 | 129460984 | 73.30% | Fig. 5 and 6 |
| RAD21TEV, CTL, noTEV, R3 | YL-NuCTL4hr-R3 | 127042108 | 27164312 | 154206420 | 82.38% | Extended Fig. 15 and 16 |
| RAD21TEV, Exp, noTEV, R3 | YL-NuExp4hr-R3 | 83913259 | 67716244 | 151629503 | 55.34% | Extended Fig. 15 and 16 |
| RAD21TEV, Exp-Shrink, noTEV, R3 | YL-NuExp4hr-shrink-R3 | 115349854 | 38016835 | 153366689 | 75.21% | Extended Fig. 15 and 16 |
| RAD21TEV, CTL, TEV, R3 | YL-NuCTL4hr-Tev-R3 | 113881113 | 23092875 | 136973988 | 83.14% | Extended Fig. 15 and 16 |
| RAD21TEV, Exp, TEV, R3 | YL-NuExp4hr-Tev-R3 | 69959740 | 64092753 | 134052493 | 52.19% | Extended Fig. 15 and 16 |
| RAD21TEV, Exp-Shrink, TEV, R3 | YL-NuExp4hr-shrink-Tev-R3 | 110465197 | 39036258 | 149501455 | 73.89% | Extended Fig. 15 and 16 |

**Extended Data Table 2. All the Hi-C libraries generated in this study.**

| ChIP experiments | ChIP libraries | Replicate | Related Figures |
|---|---|---|---|
| Nuclei in NB, IgG | YL-NB-IgG-R2 | 1 | Fig. 4h, Extended Fig. 7b |
| Nuclei in NB, CTCF | YL-NB-CTCF-R2 | 1 | Fig. 4h, Extended Fig. 7b |
| Nuclei in NB, nRAD21 | YL-NB-nRAD21-R2 | 1 | Fig. 4h, Extended Fig. 7b |
| Nuclei in NB + TEV, IgG | YL-NB-TEV-IgG-R2 | 1 | Fig. 4h, Extended Fig. 7b |
| Nuclei in NB + TEV, CTCF | YL-NB-TEV-CTCF-R2 | 1 | Fig. 4h, Extended Fig. 7b |
| Nuclei in NB + TEV, nRAD21 | YL-NB-TEV-nRAD21-R2 | 1 | Fig. 4h, Extended Fig. 7b |
| Nuclei in NBS1, IgG | YL-NBS1-IgG-R2 | 1 | Fig. 4h, Extended Fig. 7b |
| Nuclei in NBS1, CTCF | YL-NBS1-CTCF-R2 | 1 | Fig. 4h, Extended Fig. 7b |
| Nuclei in NBS1, nRAD21 | YL-NBS1-nRAD21-R2 | 1 | Fig. 4h, Extended Fig. 7b |
| Nuclei in NBS1 + TEV, IgG | YL-NBS1-TEV-IgG-R2 | 1 | Fig. 4h, Extended Fig. 7b |
| Nuclei in NBS1+ TEV, CTCF | YL-NBS1-TEV-CTCF-R2 | 1 | Fig. 4h, Extended Fig. 7b |
| Nuclei in NB + TEV, nRAD21 | YL-NBS1-TEV-nRAD21-R2 | 1 | Fig. 4h, Extended Fig. 7b |
| Nuclei in NB, IgG | YL-NB-IgG-R2 | 2 | Extended Fig. 7a and 7c |
| Nuclei in NB, CTCF | YL-NB-CTCF-R2 | 2 | Extended Fig. 7a and 7c |
| Nuclei in NB, nRAD21 | YL-NB-nRAD21-R2 | 2 | Extended Fig. 7a and 7c |
| Nuclei in NB, cRAD21 | YL-NB-cRAD21-R2 | 2 | Extended Fig. 7a and 7c |
| Nuclei in NB, IgG | YL-NB-TEV-IgG-R2 | 2 | Extended Fig. 7a and 7c |
| Nuclei in NB, CTCF | YL-NB-TEV-CTCF-R2 | 2 | Extended Fig. 7a and 7c |
| Nuclei in NB, nRAD21 | YL-NB-TEV-nRAD21-R2 | 2 | Extended Fig. 7a and 7c |
| Nuclei in NB, cRAD21 | YL-NB-TEV-cRAD21-R2 | 2 | Extended Fig. 7a and 7c |
| Nuclei in NBS1, IgG | YL-NBS1-IgG-R2 | 2 | Extended Fig. 7a and 7c |
| Nuclei in NBS1, CTCF | YL-NBS1-CTCF-R2 | 2 | Extended Fig. 7a and 7c |
| Nuclei in NBS1, nRAD21 | YL-NBS1-nRAD21-R2 | 2 | Extended Fig. 7a and 7c |
| Nuclei in NBS1, cRAD21 | YL-NBS1-cRAD21-R2 | 2 | Extended Fig. 7a and 7c |
| Nuclei in NBS1, IgG | YL-NBS1-TEV-IgG-R2 | 2 | Extended Fig. 7a and 7c |
| Nuclei in NBS1, CTCF | YL-NBS1-TEV-CTCF-R2 | 2 | Extended Fig. 7a and 7c |
| Nuclei in NBS1, nRAD21 | YL-NBS1-TEV-nRAD21-R2 | 2 | Extended Fig. 7a and 7c |
| Nuclei in NBS1, cRAD21 | YL-NBS1-TEV-cRAD21-R2 | 2 | Extended Fig. 7a and 7c |

**Extended Data Table 3. All the ChIPseq libraries generated in this study.**
